# Nanocontainer derived from silkworm carotenoprotein for carotenoid extraction and presentation in biotechnology and biomedical applications

**DOI:** 10.1101/2022.06.28.497953

**Authors:** Nikolai N. Sluchanko, Yury B. Slonimskiy, Nikita A. Egorkin, Larisa A. Varfolomeeva, Sergey Yu. Kleymenov, Mikhail E. Minyaev, Anastasia M. Moysenovich, Evgenia Yu. Parshina, Thomas Friedrich, Eugene G. Maksimov, Konstantin M. Boyko, Vladimir O. Popov

**Affiliations:** A.N. Bach Institute of Biochemistry, Federal Research Center of Biotechnology of the Russian Academy of Sciences, 119071 Moscow, Russian Federation; N.D. Zelinsky Institute of Organic Chemistry, Russian Academy of Sciences, 47 Leninsky prosp., Moscow, 119991, Russian Federation; M.V. Lomonosov Moscow State University, Faculty of Biology, 119991 Moscow, Russian Federation; Technical University of Berlin, Institute of Chemistry PC 14, Straße des 17. Juni 135, D-10623 Berlin, Germany

**Author notes:** equal contribution (Nikolai N. Sluchanko, Yury B. Slonimskiy). corresponding author: Nikolai N. Sluchanko, A.N. Bach Institute of biochemistry, Federal Research Center of Biotechnology of the Russian Academy of Sciences, Leninsky prospect 33, building 1, 119071 Moscow, Russian Federation.

**Keywords:** Carotenoprotein, antioxidant, oligomeric state, secondary structure, SEC-MALS, carotenoid transfer, liposome

## Abstract

Found in many organisms, soluble carotenoproteins are considered as antioxidant nanocarriers for biomedical applications, although the structural basis for their carotenoid transfer function, a prerequisite for rational bioengineering, is largely unknown. We report crystal structures of the Carotenoid-Binding Protein from *Bombyx mori* (BmCBP) in apo- and zeaxanthin (ZEA)-bound forms. We use spectroscopy and calorimetry to characterize how ZEA and BmCBP mutually affect each other in the complex, identify key carotenoid-binding residues, confirm their roles by crystallography and carotenoid-binding capacity of BmCBP mutants and reconstitute BmCBP complexes with biomedically-relevant xanthophylls lutein, zeaxanthin, canthaxanthin and astaxanthin. By cost-effectively and scalably solubilizing xanthophylls from various crude herbal extracts, His-tagged BmCBP remains monomeric and forms a dynamic nanocontainer delivering carotenoids to liposomes and to other carotenoid-binding proteins, which in particular makes the Orange Carotenoid Protein, a promising optogenetic tool, photoactive. Furthermore, BmCBP(ZEA) administration stimulates fibroblast growth, which paves the way for its biomedical applications.

## Main

Carotenoids are colored hydrophobic substances with profound antioxidant properties which in many organisms perform various biological functions from coloration to photoprotection and direct reactive oxygen species (ROS)-scavenging activity. Notwithstanding the great potential of carotenoids for multiple biomedical applications, their current use is limited by insolubility and poor bioavailability. Hence, natural water-soluble carotenoid-binding proteins are very promising, but so far none of them meets all necessary criteria for bioengineering and biotechnological use: i) an established spatial structure for the complex with carotenoids, ii) confirmed carotenoid transfer function and iii) a broad carotenoid-binding repertoire including carotenoids valuable to human health.

β-Crustacyanin became one of the first carotenoproteins with its crystal structure established ^1^. It forms heterodimers binding two molecules of astaxanthin (AXT) and determines the coloration of lobster carapace ^1,2^. AXT is a valuable marine carotenoid used in food supplements, medicines and cosmetics and its binding by crustacyanin was proposed to facilitate AXT concentration in biotechnology processes ^3^. However, it is not known whether crustacyanin can bind and transfer other carotenoids.

The recently discovered AXT-binding protein AstaP from eukaryotic microalgae is a stress-inducible protein conferring passive photoprotection by accumulating AXT in the cell periphery to absorb excessive sunlight ^4^. The structure and carotenoid-binding mechanism of AstaP remains to be studied, but we have recently shown that this protein can accommodate a range of different carotenoids, which makes it especially promising for carotenoid solubilization and delivery ^5^.

Perhaps the most thoroughly studied is the Orange Carotenoid Protein (OCP), a remarkable photoswitch playing a central role in cyanobacterial photoprotection ^6^. A single bound ketocarotenoid molecule, such as echinenone (ECN) and canthaxanthin (CAN), makes OCP photoactive ^7^. Upon absorption of blue-green light, it undergoes a structural transformation from the basal orange OCP^O^ form to the light-adapted red OCP^R^ form, which is capable of quenching cyanobacterial light-harvesting complexes ^6^. In addition, OCP is a good ROS quencher *in vitro* ^8^. OCP is composed of an N-terminal (NTD) and a C-terminal domain (CTD) sharing the carotenoid and being connected by an unstructured linker ^9^. Studies on separate NTD and CTD as individual proteins allowed us to discover in 2017 a unique protein-to-protein carotenoid transfer mechanism ^10,11^. Both NTD and CTD of OCP have natural homologs encoding independent carotenoproteins with confirmed roles in carotenoid transfer ^12–14^. We have shown that a CTD homolog (CTDH) can temporarily arrest the ketocarotenoid of OCP upon photoactivation of the latter, which served as proof-of-principle that water-soluble carotenoproteins can be used for the light-controlled carotenoid release ^13^. Furthermore, we demonstrated that CTDH is a potent carotenoid solubilizer and delivery module transferring carotenoids to other proteins and to membranes, which in particular alleviated oxidative stress challenge in mammalian cells ^15^. However, CTDH forms dimers, is very specific in binding ketocarotenoids and can only deliver ECN ^15^, which is not reportedly present in humans and possibly presents limited practical value. In addition, CTDH could be crystallized and structurally studied only in the apoform ^16^, which leaves its carotenoid-binding mechanism enigmatic so far.

Carotenoid-binding proteins were also found among eukaryotic proteins of the steroidogenic acute regulatory lipid transfer (START) family ^17^. A START member dubbed STARD3 was identified as the major lutein-binding protein in the human retina, where it promotes the accumulation of carotenoids (up to 1 mM) in the macula lutea ^18^. The macular carotenoids ^19^, primarily lutein (LUT) and zeaxanthin (ZEA), play a role as blue light filter and as antioxidants, decreasing the risk of age-related macular degeneration, which is one of the leading causes of blindness in the world ^20^. Despite comprehensive efforts, human STARD3 could be crystallized only in the apoform ^21,22^, which left the question of the carotenoid-binding mechanism by START domains unresolved.

Interestingly, the carotenoid-binding function of STARD3 was proposed in 2011 ^18^ based on its homology with the Carotenoid-Binding Protein from silkworm *Bombyx mori* (BmCBP) identified in 2002 ^23^. Based on 29% sequence identity to human STARD3, BmCBP was ascribed to the START protein family ^23^. BmCBP binds and transports LUT in the midgut and silk gland of silkworm larvae, which confers yellow color to the cocoons used in sericulture for thousands of years ^24^. Transposon-associated BmCBP gene inactivation or silencing yielded white cocoons and colorless hemolymph, whereas transformation of the BmCBP gene into the colorless *B. mori* strain turns cocoons and hemolymph back to yellow ^24^. BmCBP binds LUT *in vitro* and *in vivo* ^23^. Nevertheless, the structure, carotenoid-binding mechanism, ligand specificity and other properties of BmCBP remained unknown.

Here, we present high-resolution crystal structures of BmCBP in its apo- and ZEA-bound forms and describe the ZEA binding mechanism to BmCBP. We reconstitute BmCBP complexes with keto- and hydroxy-carotenoids valuable to human health and show cost-effective and scalable carotenoid enrichment by BmCBP apoprotein from various crude extracts. We define structure-activity relationships for BmCBP by crystallizing a set of its mutants and interrogating their carotenoid-binding capacity and demonstrate proof-of-principle for BmCBP applicability as antioxidant nanocarrier in different model systems.

## Results and discussion

### 1. Crystal structure of the BmCBP apoform

For our study, we engineered naïve BmCBP (297 a.a.) by removing first 67 residues to leave only its tentative ligand-binding domain and adding a cleavable His-tag on its new N-terminus to facilitate handling ^25^. The BmCBP apoform crystal structure (Supplementary Table 1 and 2) revealed in the asymmetric unit a single START-like domain at a near atomic resolution (Fig. 1a, Supplementary Fig. 1, 2a), containing unusually long Ω2-loop and β8/9 hairpin. The 12-residue long ligand-exchange Ω1-loop ^26–28^ forms a lid attached to the longest α-helix, α4, by a hydrophobic interface (Supplementary Fig. 2a). This lid covering the hydrophobic ligand-binding cavity is composed of two sides, one flexible (TAGGGR^173^G) and another rather rigid and hydrophobic (IITPR^179^). While the latter faces hydrophobic residues of the α4 helix and Ω4-loop, the flexible side of the loop is uniquely fixed by a salt bridge between its Arg173 and Asp279 of the α4 helix (Supplementary Fig. 2a,b). The electron density for Trp232 inside the unoccupied cavity is substantially dispersed, suggesting a continuum of conformations (Supplementary Fig. 2c). The deepest part of the cavity features a polar side chain of Ser206 and a salt bridge between Arg185 and Asp162 (Supplementary Fig. 2c).

**Fig. 1.**
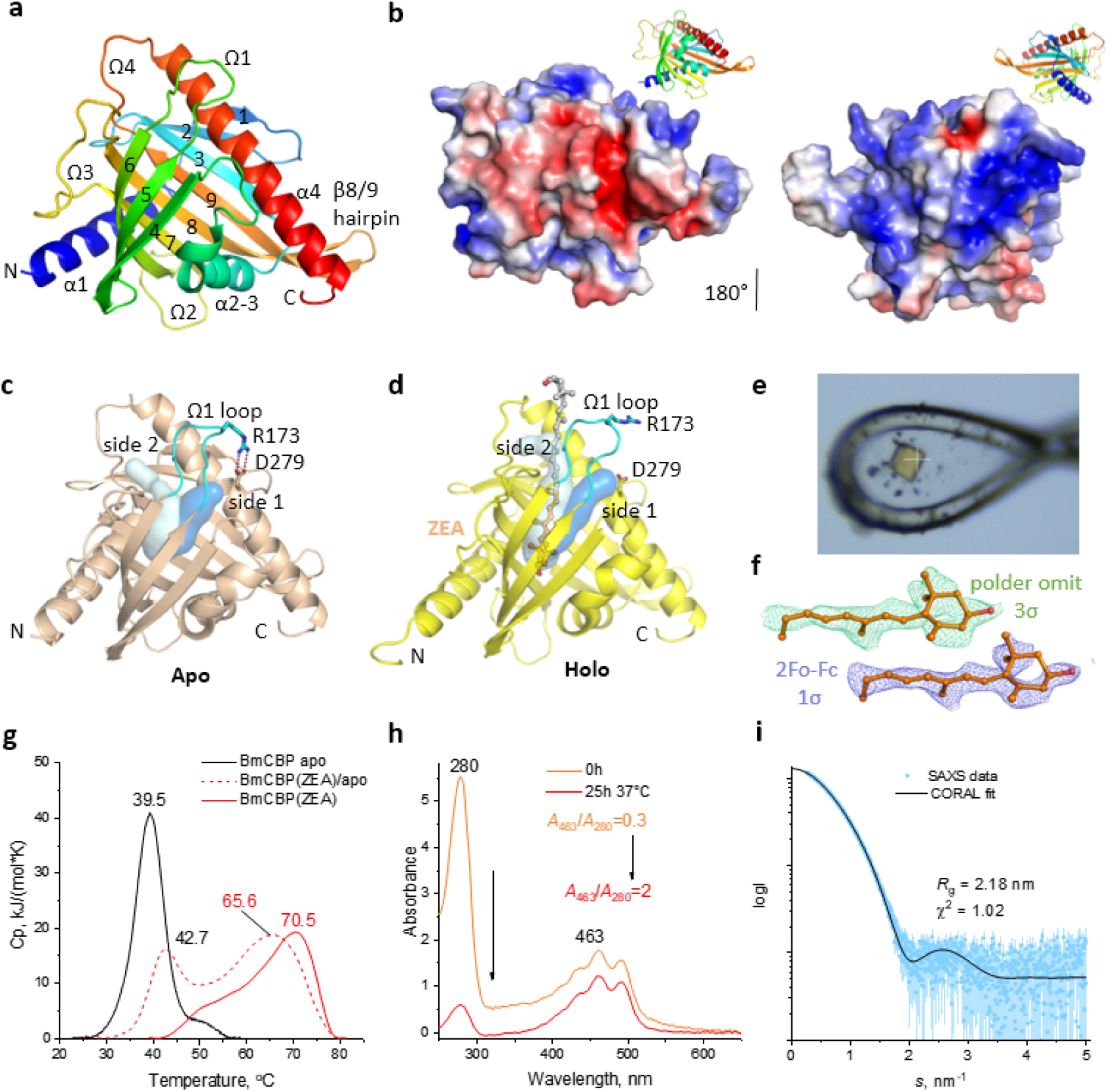
Apo and zeaxanthin-bound BmCBP. a. Overall view showing BmCBP apoprotein as a ribbon diagram colored by a gradient from the N (blue) to C terminus (red). α1-α4 helices and Ω1-Ω4-loops are labeled, β-strands are numbered 1-9. b. Opposite sides of the BmCBP surface reveal distinct patterns of the electrostatic potential colored by a gradient from red (−3 kT/e) to blue (+3 kT/e). For clarity the views are doubled as ribbon diagrams (color gradient from N to C terminus) in the top right corners. 1.45-Å crystal structure of the apoform (c) and 2.0-Å crystal structure of ZEA-bound BmCBP (d) showing by cyan and blue semi-transparent surfaces tentative carotenoid-binding tunnels computed by CAVER ^29^ (a 1.1-Å minimum probe radius). For the holoform, ZEA was removed from the model before computation. e. A microphotograph showing the yellow BmCBP(ZEA) crystal during X-ray data collection. The white cross has a 30-μm size. f. The ZEA portion supported by the polder omit ^30^ and 2Fo-Fc maps. Contour levels are indicated. g. DSC thermograms for the BmCBP apoform, holoform, and an apo-/holo-mixture heated at a 1 °C/min rate. *T*_m_ values for the peaks are shown in °C. h. Effect of selective melting of BmCBP apoform on the absorbance spectrum of the apo/holo mixture. i. The fit of the SAXS data in solution by the crystal structure of BmCBP(ZEA) supplied with the missing N-terminal residues.

The BmCBP exterior has a remarkable distribution of opposite electrostatic surface potentials grouped on two sides of the globule (Fig. 1b), which may be related to the protein function. According to CAVER ^29^, BmCBP has two major tunnels potentially suitable for the ligand entry into the cavity with the “gates” located at two sides of the Ω1-loop (side 1 and 2, Fig. 1c). While side 2 is more hydrophobic and thus more complementary to the lipidic ligands, side 1 features the Arg173-Asp279 salt bridge, a potential obstacle for ligand entry. On side 2, at the base of the Ω1-loop resides the gatekeeper Arg179 (Supplementary Fig. 2b), which H-bonds to Thr177 and the carbonyl backbone of the Ω1-loop. While in our structure it is diverted, when unleashed, the Arg179 side chain may control access to the ligand-binding cavity.

### 2. Structural insights into carotenoid binding by BmCBP

Further, we solved the crystal structure of BmCBP holoprotein reconstituted in a ZEA-synthesizing *E.coli* strain ^25^ (Fig. 1d and Supplementary Tables 1, 2). The presence of ZEA in BmCBP was strictly verified by spectrochromatograms (see below) and the intense yellow color of the crystals (Fig. 1e). Scrupulous refinement of the protein and solvent modeling were required to reveal a rather weak but continuous electron density inside the BmCBP cavity attributable to parts of a ZEA molecule. This was sufficient to build 20 atoms of ZEA including its β-ring and part of the polyene chain into the density for further structure refinement. While the final refined structure still had density only for a part of the carotenoid (Fig. 1f), the position of the inner β-ring and the direction of the polyene chain allowed extending the carotenoid model beyond the region supported by electron density (Fig. 1d). According to such modeling, a ~30 Å long ZEA molecule inevitably protrudes from the cavity, exiting at side 2 of the Ω1-loop. In our model, the carotenoid occupies the rather straight tunnel decorated by hydrophobic side chains (Fig. 1d), whereas exit at side 1 would likely require *trans/cis* isomerization beyond the region supported by electron density. Such a configuration would yield a clear near-UV absorbance peak at 330-350 nm ^31,32^; however, such a band is not observed in the absorbance spectra of BmCBP-carotenoid complexes (see below).

The BmCBP(ZEA) structure reveals a rearrangement of the Ω1-loop at side 1 involving a disruption of the salt bridge between R173 and D279, which became >8 Å apart (Fig. 1d). Such an Ω1-loop rearrangement can result from the carotenoid pushing the loop from the hydrophobic side 2 (Cα atoms of I175 and I176 are displaced by 0.7 and 0.5 Å, respectively, relative to their positions in the apoform) and thereby deforming the flexible side 1 (Cα atoms of R173, G172 and G171 are displaced by 2.1, 6.2 and 1.5 Å, respectively). Upon embedment in the tunnel, ZEA likely undergoes different bending distortions, which disperses the electron density for the carotenoid in the outer region.

Overall, the BmCBP backbone did not change much upon ZEA binding (Cα RMSD=0.32 Å for 204 out of 238 aligned atoms, Supplementary Fig. 1). Nevertheless, differential scanning calorimetry indicated that ZEA binding dramatically (by ~30 °C) stabilizes BmCBP (*T*_m_ increases from 39.5 to 70.5 °C; Fig. 1g). The denaturation profile of a BmCBP(ZEA) preparation having a *A*_462_/*A*_280_ ratio of ~1.3 and containing ~50% of the apoform, showed two incompletely resolved peaks (*T*_m_ = 42.7 °C and 65.6 °C, respectively) (Fig. 1g). While the first peak had a ~3 °C higher *T*_m_, its shape and position were similar to that of the individual apoform. The second, broad peak with the higher *T*_m_ is missing from the thermogram of the apoform and mostly represents melting of the BmCBP(ZEA) complex, yet its *T*_m_ is lower than that for the pure BmCBP(ZEA) (65.6 vs 70.5 °C). These observations likely reflect transient nature of BmCBP-carotenoid complexes between which carotenoids are constantly exchanged by the excess of BmCBP molecules during heating, which apparently stabilizes the apoform proportion and destabilizes its holoform counterpart. In the ligand absence, BmCBP is a rather unstable protein whose thermal transition starts already at 30 °C. Based on this finding, we demonstrated the possibility of selectively melting the apoprotein via continuous incubation at 37 °C, which after centrifugation made BmCBP holoprotein very pure (Fig. 1h). We propose that this simple procedure could be used to make clean BmCBP holoprotein preparations in large scale.

The protrusion of the BmCBP-embedded carotenoid (Fig. 1d) likely provides a lever for carotenoid release to other proteins but is insufficient for a second BmCBP monomer to form a carotenoid-stabilized homodimer, like in cyanobacterial CTDH carotenoproteins ^13^. In agreement, small-angle X-ray scattering (SAXS) confirmed that the crystal structure of a single BmCBP(ZEA) molecule adequately describes the conformation in solution (Fig. 1i). First, the SAXS-derived *M*_w_ values directly confirmed the monomeric status of the protein (Supplementary Table S3). Second, the CORAL-derived ^33^ model (*R*_g_=2.18 nm, experimental *R*_g_=2.12 nm, *D*_max_=8.5 nm) provided an exceptionally good approximation of the experimental SAXS profile (*X*^2^=1.02). Thus, BmCBP forms a unique carotenoid-embedding nanocontainer not requiring dimerization to accommodate its ligand.

In the crystal structure, ZEA lies within the hydrophobic tunnel so that the hydroxyl group at the inner β-ring forms polar contacts with Ser206 (2.6 Å) and one of the alternative conformations of Trp232 (3.5 Å) (Fig. 2a). The distance from the hydroxyl oxygen of the carotenoid to the side chain nitrogen of Arg185 is likely too large to form a stable polar contact (4.0 Å). The side chain of this arginine, although rather flexible on its own, in the BmCBP structure is firmly kept out of reach of the carotenoid ring by several H-bonding interactions with the backbone carbonyls of residues Phe134 and Lys135, and the Asp162 side chain (Fig. 2a). The inner β-ring stacks with both Trp232 rotamers located in a plane at ~4 Å distance from the plane of the carotenoid ring. One Trp232 rotamer places the indolyl nitrogen at a distance sufficient for a weak H-bond with the carotenoid hydroxyl (3.5 Å), albeit at a suboptimal angle. The polder omit density map ^30^ for the ligand directs the polyene chain along the hydrophobic tunnel (Fig. 2a), but the embedded carotenoid conformation likely differs from a straight line.

**Fig. 2.**
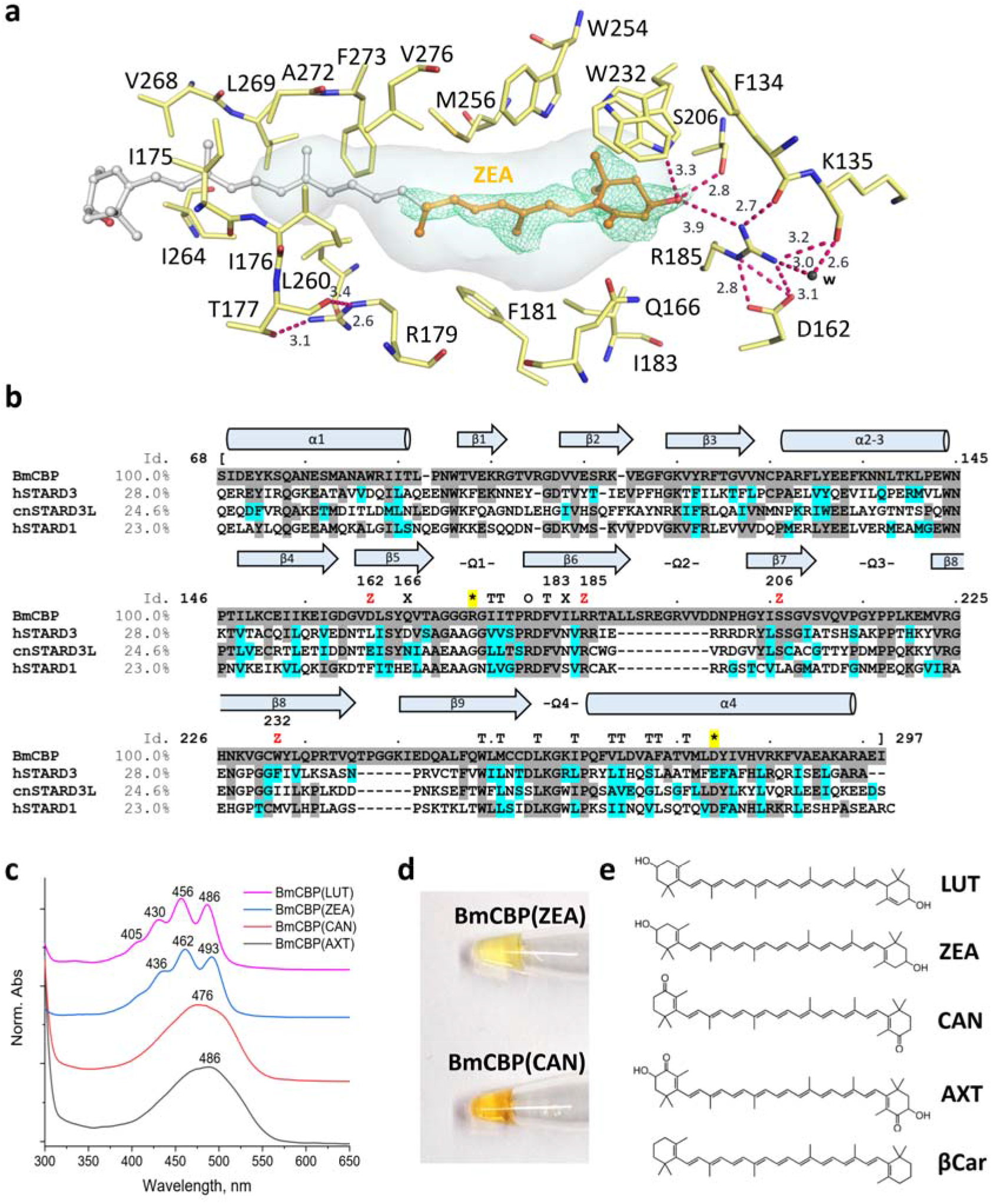
Structural determinants of carotenoid binding by BmCBP’s START domain. a. Close-up view of the ligand-binding cavity of BmCBP in the crystal structure of the holoform showing the position of ZEA and pigment-protein interactions. The semi-transparent grey surface shows the tunnel computed by CAVER ^29^. A green mesh shows a polder omit map ^30^ for the ZEA fragment contoured at 3σ. Key residues involved in carotenoid binding and forming the carotenoid-binding tunnel, as well as several characteristic distances (in Å) are indicated. b. Aligned START domains of BmCBP (Uniprot Q8MYA9), hSTARD3 (Uniprot Q14849), hSTARD1 (Uniprot P49675) and cnSTARD3L (Uniprot A0A0C5B5I7). Residues identical to those in BmCBP are highlighted by grey, similar residues are highlighted by cyan according to the five similarity groups: D,E / W,Y,F / R,K,H / N,Q / M,V,I,L,A / G,C,P,S,T. Secondary structures and key loops corresponding to those in the BmCBP crystal structure are depicted above the alignment. “T” indicates residues forming the carotenoid tunnel, “Z” indicates residues surrounding or contacting the carotenoid ring in the crystal structure. “O” marks the conserved gatekeeper arginine at the base of the Ω1-loop. Asterisks highlighted in yellow denote the Arg173-Asp279 salt bridge immobilizing the Ω1 loop on the α4-helix. c. Normalized absorbance spectra of BmCBP complexes with different carotenoid types as recorded by spectrochromatography. d. Color of the purified BmCBP holoforms with ZEA and CAN reconstituted in the corresponding carotenoid-producing *E.coli* cells. e. Structural formulae of lutein (LUT), zeaxanthin (ZEA), canthaxanthin (CAN), astaxanthin (AXT) and β-carotene (βCar).

Many residues in the carotenoid-binding tunnel of BmCBP are identical or similar in human STARD3, another recently reported carotenoid-binding STARD3-like protein from golden scallop *Chlamys nobilis* (cnSTARD3L) ^34^, and human STARD1 (Fig. 2b). Ser206 coordinating the carotenoid hydroxyl in BmCBP is also present in hSTARD3 and cnSTARD3, all of which are reported carotenoid binders ^18,23,34^, but is replaced by Leu199 in hSTARD1, for which carotenoid binding is not documented. Therefore, this serine likely plays a role in coordinating hydroxylated carotenoids. While Arg185 is invariably present in all homologs analyzed (Fig. 2b), its neighbor Asp162 becomes either Leu328 in hSTARD3 or Phe165 in hSTARD1, and only in cnSTARD3L is replaced by a synonymous Glu. One can expect that elimination of the negative charge at this position would unleash conformational mobility of conserved Arg185, potentially affecting ligand binding. The roles of other variable residues in positions 166, 183 and 232 of BmCBP are less obvious.

We have reconstituted BmCBP complexes with ZEA and its isomer LUT, which differs by the position of one double bond in its ε-ring ^25^. Based on our structural data, we suggested that the ligand specificity of BmCBP is rather broad and reconstituted its holoforms with a range of carotenoids (Fig. 2c,d). In principle, not only 3,3’-hydroxylated xanthophylls ZEA and LUT, but also those containing either only 4,4’-ketogroups (CAN) or simultaneously 4,4’-ketogroups and 3,3’-hydroxyl groups (AXT) could be embedded into BmCBP, each producing distinct spectral signatures (Fig. 2c). All our attempts to reconstitute BmCBP complexes with βCar were unsuccessful, possibly because this carotenoid lacks polar groups, which may be critical for the carotenoid uptake. The principle ability of BmCBP to bind LUT, ZEA, CAN and AXT (Fig. 2e), all having chemically different carotenoid rings, suggested that the hydrophobic tunnel provides BmCBP with the unique ability to accommodate various lipophilic antioxidants.

We next analyzed BmCBP ability to capture carotenoids from various natural sources differing by the carotenoid content (Fig. 3a). Interestingly, BmCBP apoprotein exhibited strong ability to enrich carotenoids from crude spinach methanolic extracts that initially contained numerous pigments (Fig. 3b). This was further confirmed by spectrochromatography. As a reference, BmCBP apoprotein efficiently extracted ZEA and LUT present in commercial food supplements (Fig. 3c), yielding the absorbance spectrum with the maxima positions intermediate between those for LUT or ZEA individual spectra (Fig. 3d and 2c). Likewise, BmCBP efficiently extracted carotenoids from spinach, dandelion, parsley and calendula extracts, all containing xanthophylls (their different relative content is reflected in different spectral shifts and shapes, Fig. 3e), but not from the carrot extract, where a prevalent carotenoid is known to be βCar. This agrees with our data on reconstitution of BmCBP complexes with individual carotenoids (Fig. 2). Thus, BmCBP apoprotein can be loaded with natural xanthophylls present in crude extracts, and this cost-effective one-step procedure enables the pronounced carotenoid enrichment compatible with biotechnological scale-up.

**Fig. 3.**
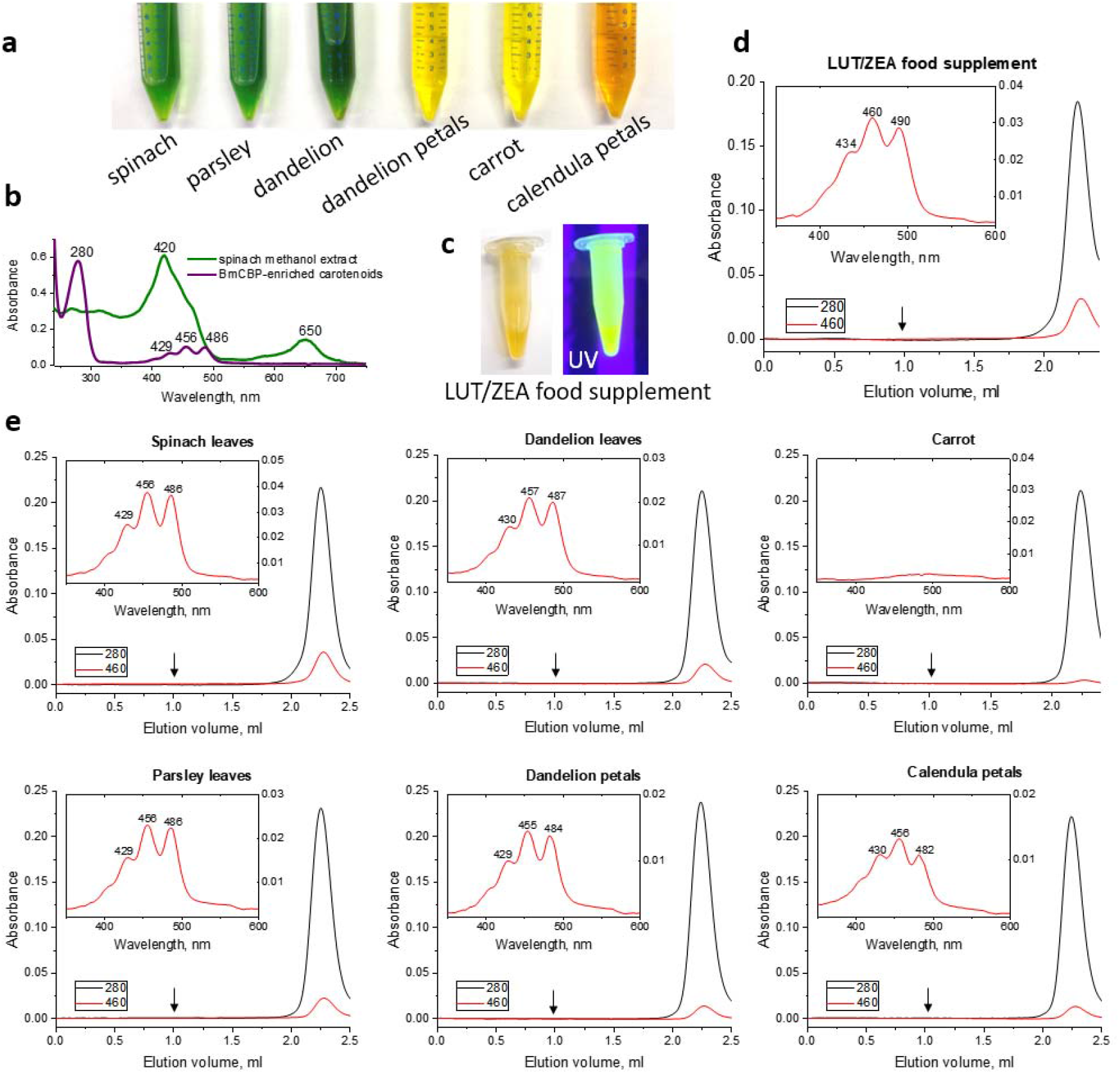
BmCBP apoprotein extracts carotenoids from natural sources. a, Tubes with herbal methanolic extracts. b, Extraction and enrichment of the carotenoid fraction from spinach using BmCBP apoprotein. c, An Eppendorf tube with the methanolic extract of Ocuvite^®^ food supplement containing LUT and ZEA and used as a reference. d,e. Analysis of BmCBP-mediated carotenoid extraction from different sources by SEC with full-spectrum detection. SEC profiles are shown with the absorbance spectra in the inserts. Arrows indicate column void volume. d, Reference BmCBP-mediated extraction of xanthophylls from a LUT/ZEA-containing food supplement. e, BmCBP-mediated extraction of carotenoids from herbs shown on panel a. The main absorbance maxima are indicated in nm.

### 3. BmCBP binding changes optical characteristics of the carotenoid

Spectral properties of carotenoids are highly sensitive to their conformation and microenvironment. The shape of the carotenoid absorption spectrum is determined by a combination of electronic and vibrational transitions, which may give a pronounced fine structure. The effective size of the conjugated π-electron system depends on the number of double bonds in plane and determines the position of the absorption maximum and the Raman shifts, which are interrelated ^35^. The maximum possible effective conjugation length of ZEA is one double bond longer than that of LUT because of the arrangement of double bonds in ε and β-ionone rings (Fig. 3e). This explains the blue-shifted absorption and a greater Raman shift of the C=C double bond oscillation (v_1_ of LUT than of ZEA in methanol (Fig. 4a,b). The equilibrium carotenoid conformation in methanol has a ~40-50° dihedral angle between the ionone ring and polyene ^36^, which does not allow it to reach the maximum effective conjugation length. However, carotenoid embedment in BmCBP produces a bathochromic absorption shift and lowers the v_1_ frequency (Fig. 4a,b), therefore, the effective conjugation length increases; this indicates an altered carotenoid conformation and partial immobilization of the carotenoid ring. Since the double bond of the ε-ring of LUT is not conjugated with the polyene chain, we assume that it is the β-ring that forms contacts with the protein, which increases the conjugated length. In ZEA, both rings are chemically equivalent and have no apparent preferences for entering the BmCBP cavity. Similar v_1_ frequencies for LUT and ZEA in BmCBP may indicate their binding conformation is similar, in which a β-ring is specifically oriented in the protein while the second ring (either ε or β) protrudes to the solvent.

**Fig. 4.**
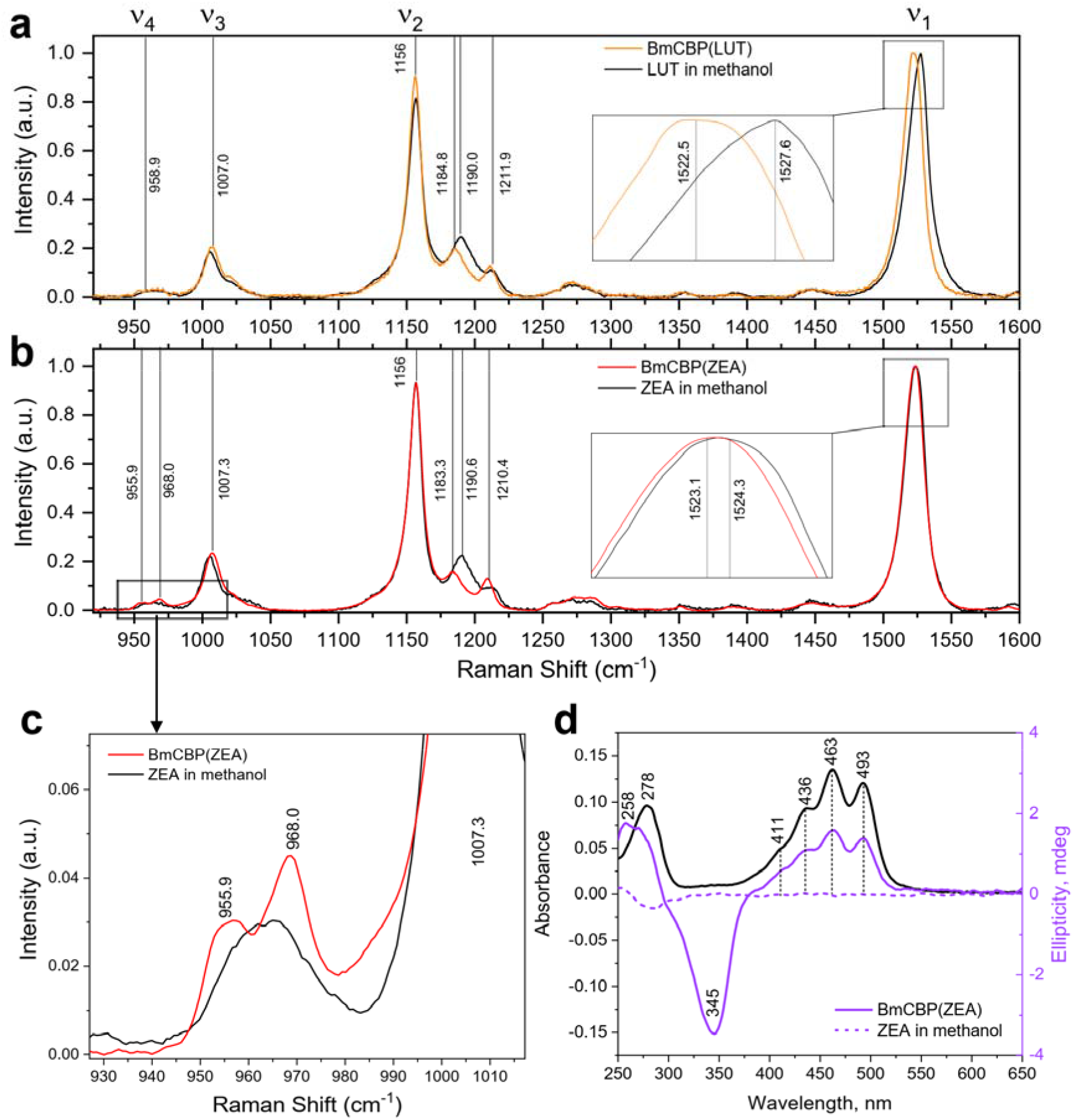
BmCBP changes spectral properties of the bound carotenoid. a. Raman spectra of LUT in BmCBP and in methanol. A magnified view of the v_1_ band is shown as an insert. b. Raman spectra of ZEA in BmCBP and in methanol. A magnified view of the v_1_ band is shown as an insert. c. A magnified view of the v _4_ region of the spectrum presented in panel b showing the HOOP peak at 968 cm^-1^. d. Visible CD (magenta) and absorbance (black) spectra of ZEA in BmCBP-bound form. Vis-CD spectrum of ZEA in methanol (dashed magenta) is shown for comparison.

The appearance of the peak at 968 cm^-1^ accompanying ZEA binding to BmCBP (Fig. 4c), associated with the so-called hydrogen-out-of-plane (HOOP) wagging mode ^37,38^, strongly indicates torsional bending of the ZEA polyene. Such anisotropy is further supported by the induced circular dichroism: unlike the strictly symmetric ZEA in methanol, ZEA embedded in BmCBP has a pronounced chirality (Fig. 4d). Therefore, BmCBP binding not only limits the conformational mobility of the carotenoid, as indicated by the fine structure of the absorbance spectrum, but also forces polyene bending. Since upon ZEA binding to BmCBP the peak at 968 cm^-1^ reaches only ~20% of the v_3_ peak intensity (Fig. 4b), the ZEA bending curvature in BmCBP is likely less pronounced than in the case of the ketocarotenoid in OCP, for which the amplitude of the HOOP peak reaches ~80% of the v_3_ peak ^37^, and the curvature has a 28 Å radius and ~16° deviation from all-trans conformation (180°) ^9^. This is in line with our structural data.

### 4. Structure-activity relationships in BmCBP with the mutated carotenoid-binding site

Although hSTARD3 was reported to bind several carotenoids including ZEA *in vitro* ^18^, all our attempts to reconstitute its complexes with any carotenoid were not successful (see Supplementary Fig. 3). Focusing on the detected differences in the organization of the tentative carotenoid-binding site between the two proteins (see Supplementary Fig. 3), we engineered single BmCBP mutants where BmCBP residues were replaced by those of hSTARD3, i.e. D162L, W232F, Q166D and I183N. Since the semi-conserved Ser206 directly contacts the hydroxyl of the carotenoid in BmCBP (Fig. 2a), we also designed the S206V mutant as a control. We determined crystal structures of D162L, W232F and S206V apoproteins (Supplementary Tables 1,2 and Supplementary Fig. 4) and used all above-mentioned mutants to assess their carotenoid-binding capacity (Fig. 5).

**Fig. 5.**
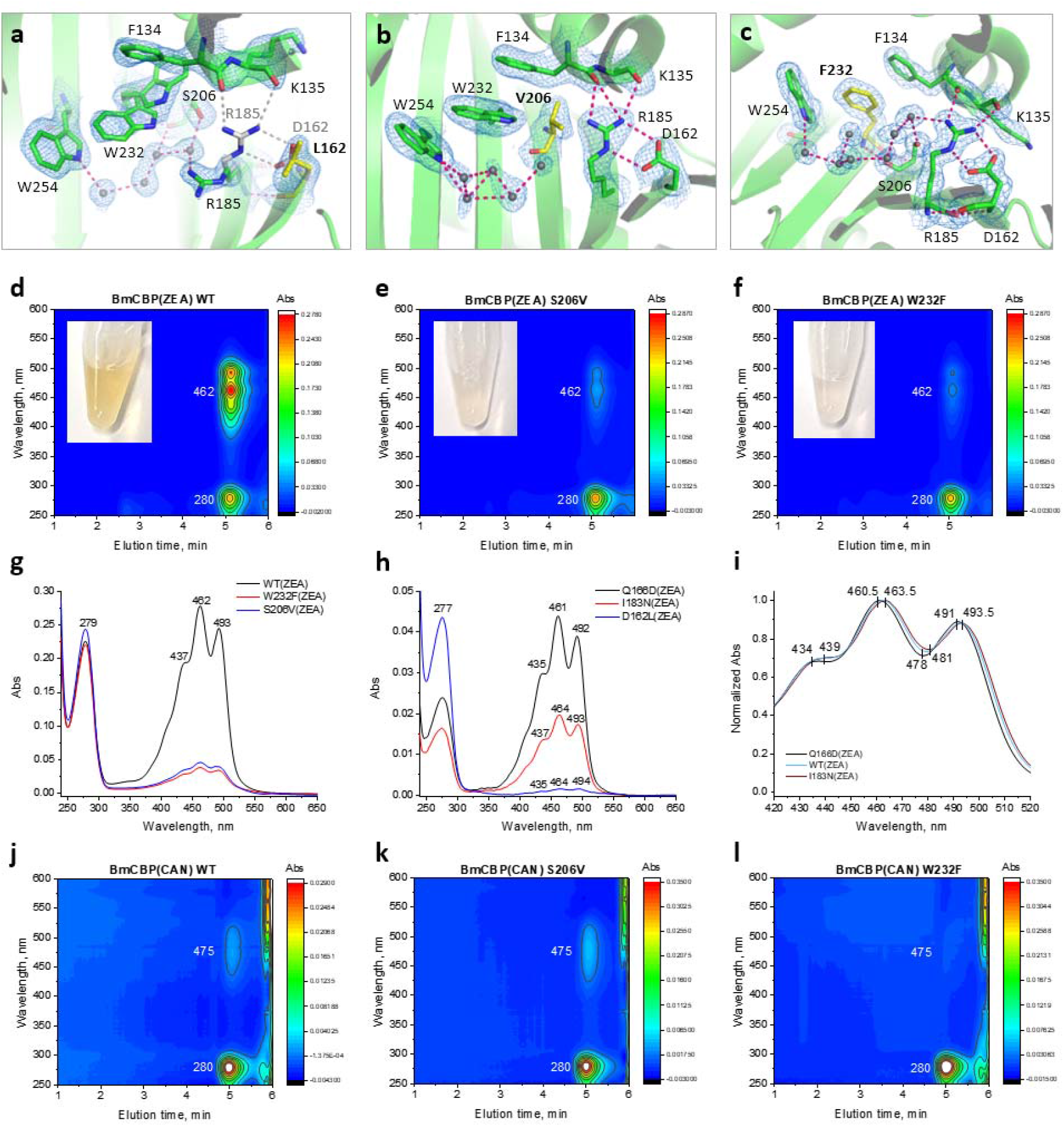
Crystal structures and carotenoid-binding capacity of the engineered BmCBP mutants. a-c. Magnified views of the carotenoid-binding site of the S206V (b), W232F (c) and D162L (d) mutants validating the mutations introduced (in yellow). In a, grey sticks show the position of the R185 and D162 residues forming the salt bridge in BmCBP WT. The mutation led to the disruption of the salt bridge and R185 displacement towards the ligand-binding cavity. The 2Fo-Fc electron density maps contoured at 1σ are demonstrated. d-l. The His-tagged proteins were expressed in ZEA-producing *E.coli* cells, purified by IMAC and subjected to spectrochromatography (Superdex 200 Increase 5/150, 0.45 ml/min). d-f. Spectrochromatograms showing the purity and typical UV-Vis absorbance of the WT (d), S206V (e) or W232F (f) variants. The inserts show the color of the samples loaded on the column. Unlike yellow WT, the mutants have dramatically reduced Vis absorbance. g. Absorbance spectra of the BmCBP variants presented in d-f. h. Absorbance spectra of the other three BmCBP variants obtained at reduced yields and additionally concentrated prior to spectrochromatography. i. A magnified view showing slight spectral differences in the carotenoid absorbance of the selected BmCBP variants. j-l. Spectrochromatograms of BmCBP holoforms with CAN for WT (j), S206V (k) and W232F (l). Note that while WT and S206V bind CAN comparably, W232F shows no absorbance in the visible region and hence no CAN binding.

Similarity of new crystal structures (Cα RMSD <0.45 Å when superimposed with the BmCBP WT apoprotein structure, Supplementary Fig. 4a), indicating unaltered protein folding. In all structures, the mutated residues were clearly supported by the electron density (Fig. 5a-c). In S206V and W232F structures, we did not find other substantial changes, whereas in the D162L structure, the mutation disrupted the salt bridge with Arg185, as expected. As a result, H-bonds with the backbone carbonyls of Phe134 and Lys135 also broke, which displaced the Arg185 side chain to the ligand-binding cavity, much like Arg351 in hSTARD3 (Supplementary Fig. 3d).

After confirming the expected modifications of the carotenoid-binding site of BmCBP by crystallography, we assessed the ability of all designed BmCBP mutants to form complexes with ZEA upon expression in ZEA-synthesizing *E. coli* cells, with the wild-type BmCBP used as a positive control (Fig. 5d). Despite similar expression levels of all mutants (Supplementary Fig. 4b), only the WT, W232F and S206V variants could be purified in large amounts and analyzed by spectrochromatography directly after IMAC (Fig. 5d-g). For Q166D, I183N and especially D162L, a low yield of the soluble protein required additional purification by SEC and concentration prior to spectrochromatography (Fig. 5h). For all BmCBP variants, we compared the absorbance spectra from their peaks on spectrochromatograms (Fig. 5g,h). In a striking contrast with WT, the S206V and W232F mutants displayed substantially reduced Vis/UV absorbance ratios and hence carotenoid-binding capacities (Fig. 5g). While the D162L mutant was expressed at a low yield, its absorbance spectrum contained only minor peak characteristic for ZEA compared with the UV peak (Fig. 5h), which indicated low stability of this protein and its poor carotenoid-binding capacity. These data strongly supported the functional relevance of Ser206, Trp232 and Asp162 residues in ZEA binding. Notably, hSTARD1, despite having a similar fold (PDB 3P0L, Cα (RMSD=1.02 Å when superposed with the BmCBP apo structure), has Leu199 instead of Ser206 in BmCBP, Met225 instead of Trp232 in BmCBP and Phe165 instead of Asp162 in BmCBP. Our data predict the inability of hSTARD1 to bind ZEA.

Despite their low yield, comparable to that of the D162L mutant, the Q166D and I183N mutants could still be obtained in the holoform and showed large Vis/UV absorbance ratios indicating their principle ability to bind ZEA (Fig. 5h). This indicates that, in contrast to the inhibitory effects of the S206V, W232F and D162L substitutions, the neutral effects of the Q166D and I183N substitutions suggest that residues in these positions are probably not essential for carotenoid binding. Yet, the Q166D and I183N mutations caused subtle shifts of the absorbance in the visible region (Fig. 5i), in line with the close proximity of these residues to the carotenoid-binding site in BmCBP. This illustrates how sensitive to its microenvironment the spectral properties of the carotenoid are.

Noteworthily, we found that CAN binding efficiency is comparable for BmCBP WT and the S206V mutant (Fig. 5j,k), in which the key Ser206 side chain coordinating the ZEA hydroxyl (Fig. 2a) is replaced by the hydrophobic group of valine lacking this ability. Given that CAN has 4,4’-ketogroups instead of 3,3’-hydroxygroups of ZEA, this observation is in line with the structure of the BmCBP(ZEA) complex and is likely explained by the fact that the CAN ketogroup in BmCBP S206V points away from the 206 residue and thus does not experience an effect of the S206V substitution. Such orientation of the carotenoid ring in the BmCBP cavity is further supported by the relatively low efficiency of BmCBP WT binding to CAN than to ZEA (Fig. 5d and j), because only the ZEA hydroxyl can form contacts with Ser206. In contrast, under the same conditions, CAN binding to the W232F mutant was insignificant (Fig. 5l), which confirms the importance of the indolyl group of the 232 residue.

### 5. BmCBP is a dynamic carotenoid shuttle

After dissecting the carotenoid-binding mechanism of BmCBP, we asked if this nm-sized protein could be used for carotenoid delivery to biological membrane models and other proteins. When mixed with liposomes and then analyzed by spectrochromatography, BmCBP(ZEA) displayed an efficient carotenoid transfer accompanied by a depletion of the carotenoid-specific absorbance in the protein fraction and a concomitant appearance of the carotenoid absorbance in the liposome fraction (Fig. 6a and Supplementary Fig. 5a). After Rayleigh scattering subtraction, ZEA absorbance in liposomes revealed a much less pronounced vibronic structure than in BmCBP (Fig. 6b), reflecting a distinct chemical environment and thus physical migration of the carotenoid.

**Fig. 6.**
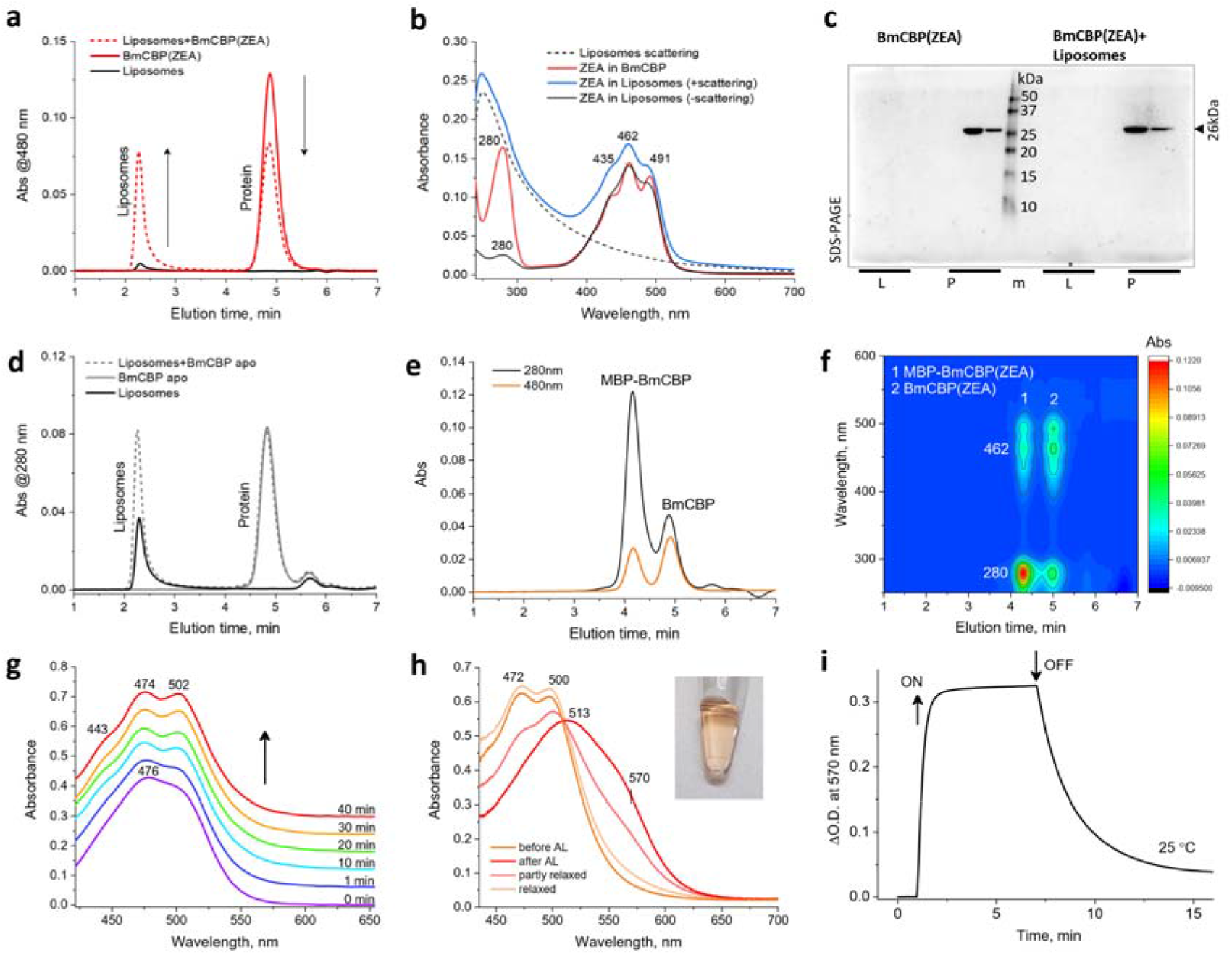
BmCBP is a dynamic carotenoid shuttle. a. ZEA transfer from BmCBP to liposomes studied by spectrochromatography (Superdex 200 Increase 5/150, 0.45 ml/min). Arrows indicate the changes of the carotenoid absorbance. b. Absorbance spectra of ZEA in BmCBP and in liposome fractions. c. Interaction of the BmCBP(ZEA) with liposomes assessed by SEC followed by SDS-PAGE of the liposome (L) and protein (P) fractions. Mass markers are shown in kDa (lane “m”). Note that BmCBP does not migrate into the liposome fraction, suggesting no stable association with the membranes. d. The BmCBP apoform does not interact with liposomes. e,f. ZEA is dynamically repartitioned between BmCBP and MBP-BmCBP as revealed by SEC profiles (e) with continuous absorbance spectrum detection (f). g. CAN transfer from BmCBP to the apoform of *Synechocystis* OCP followed by changes of absorbance. The arrow shows the time course, time after mixing is indicated. Note the evolution of BmCBP(CAN) spectral signatures into the signatures of OCP(CAN). h. The photoactivity of OCP(CAN) species formed after CAN transfer from BmCBP followed by absorbance changes upon illumination by a blue LED (445 nm). The insert shows the orange color of the sample after CAN transfer to OCP. AL, actinic light. i. The OCP photocycle in a kinetic regime monitored by absorbance at 570 nm upon switching the LED on and off (see arrows).

Noteworthily, fusion of BmCBP to the C terminus of the maltose-binding protein (MBP-BmCBP) did not abolish the ability of BmCBP to bind and transfer ZEA (Supplementary Fig. 5b,c), which is beneficial for constructing modular antioxidant delivery systems based on BmCBP in the future.

Unlike reported for the microalgal AstaP carotenoprotein ^5^, BmCBP did not display tight physical association in the holo- (Fig. 6c) and apoform (Fig. 6d) with lipid membranes as no protein was detected in the liposome fraction by SDS-PAGE. Therefore, BmCBP only transiently approaches membranes to deliver its carotenoid, which may be favored by the peculiar distribution of charges on the BmCBP surface (Fig. 1b). Assuming BmCBP is a dynamic carotenoid shuttle, we tested if carotenoids are exchanged between BmCBP molecules by mixing BmCBP(ZEA) as a donor (28 kDa) and an MBP-tagged BmCBP apoprotein (68 kDa) as an acceptor of the carotenoid. As expected, the SEC profile of such a mixture revealed two peaks with the carotenoid-specific absorbance representing MBP-BmCBP and BmCBP, respectively (Fig. 6e,f), indicating carotenoid repartitioning between the BmCBP molecules, not precluded by the MBP-tag.

Seeking for a good reporter of the successful carotenoid transfer, we used cyanobacterial OCP apoprotein as a carotenoid acceptor from BmCBP. Upon binding ketocarotenoids, the OCP apoform undergoes compaction and acquires the characteristic absorbance spectrum in the dark-adapted state, while experiencing a dramatic spectral shift upon transition to the light-adapted state, with a photocycle that can be followed by changes of absorbance at 550-570 nm ^10,11^. For this experiment, the BmCBP complex with the ketocarotenoid canthaxanthin (CAN) was used. Upon mixing it with the *Synechocystis* OCP apoform, lacking any absorbance in the visible spectral region, we observed a relatively slow (~40 min) transformation of the absorbance spectrum into the one typical for the dark-adapted OCP ^10^ (Fig. 6g). Upon completion of CAN transfer, the sample could be reversibly photoactivated by a 445-nm light-emitting diode and the typical OCP(CAN) photocycle could be recorded (Fig. 6h,i).

This confirms that BmCBP can transfer carotenoids to other proteins; the time-course of this process (minutes) is suitable for the controlled release of antioxidants by BmCBP. Our recently described genetically encoded OCP-based fluorescent thermometer ^39^ is among devices that can significantly benefit from the targeted CAN delivery from BmCBP.

### 6. BmCBP delivers zeaxanthin to fibroblasts promoting their growth

After establishing that BmCBP is capable of transferring carotenoids into model liposome membranes, enriching them with antioxidant molecules, we tested this reaction in a cell model. Being useful for studying cytotoxicity ^40^, primary mouse embryonic fibroblasts (MEF) were used to determine BmCBP biocompatibility *in vitro*. For this purpose, 0.5 μM BmCBP(ZEA) (per carotenoid) was added to cells, and similar concentrations of the apoprotein, ZEA in DMSO, or DMSO, were used as controls (Fig. 7). Apparently, BmCBP(ZEA) stimulated growth of MEFs (Fig. 7a-d). The results of the MTT-test show that cell metabolic activity remained high throughout the cultivation time, moreover, on day 7, the amount of formazan crystals significantly increased in the group of cells incubated with BmCBP(ZEA) compared to the other groups (Fig. 7e). Statistically significant differences were also found for the ZEA/control pair, which confirmed the protective function of BmCBP(ZEA) and ZEA under conditions of increased cell density and deficiency of oxygen and nutrient resources in monolayer cultures. The number of cells detected on day 7 of cultivation was similar for all groups tested, however, on day 3, the difference was significant for the BmCBP(ZEA) group and the rest of the samples (Fig. 7f). This effect is likely associated with the increased bioavailability of ZEA complexed with BmCBP and is multifaceted. Carotenoids can not only neutralize free radicals but also modulate the activity of retinoic acid receptors ^41^, which in turn play a critical role in the function of various mammalian tissues, including skin ^42^, nerve ^43^ and cardiac tissue ^44^. Given the high stability of carotenoids in water-soluble carotenoproteins and better bioavailability compared to liposomes, we assume that this method of carotenoid delivery may be in demand for various biomedical applications.

**Fig. 7.**
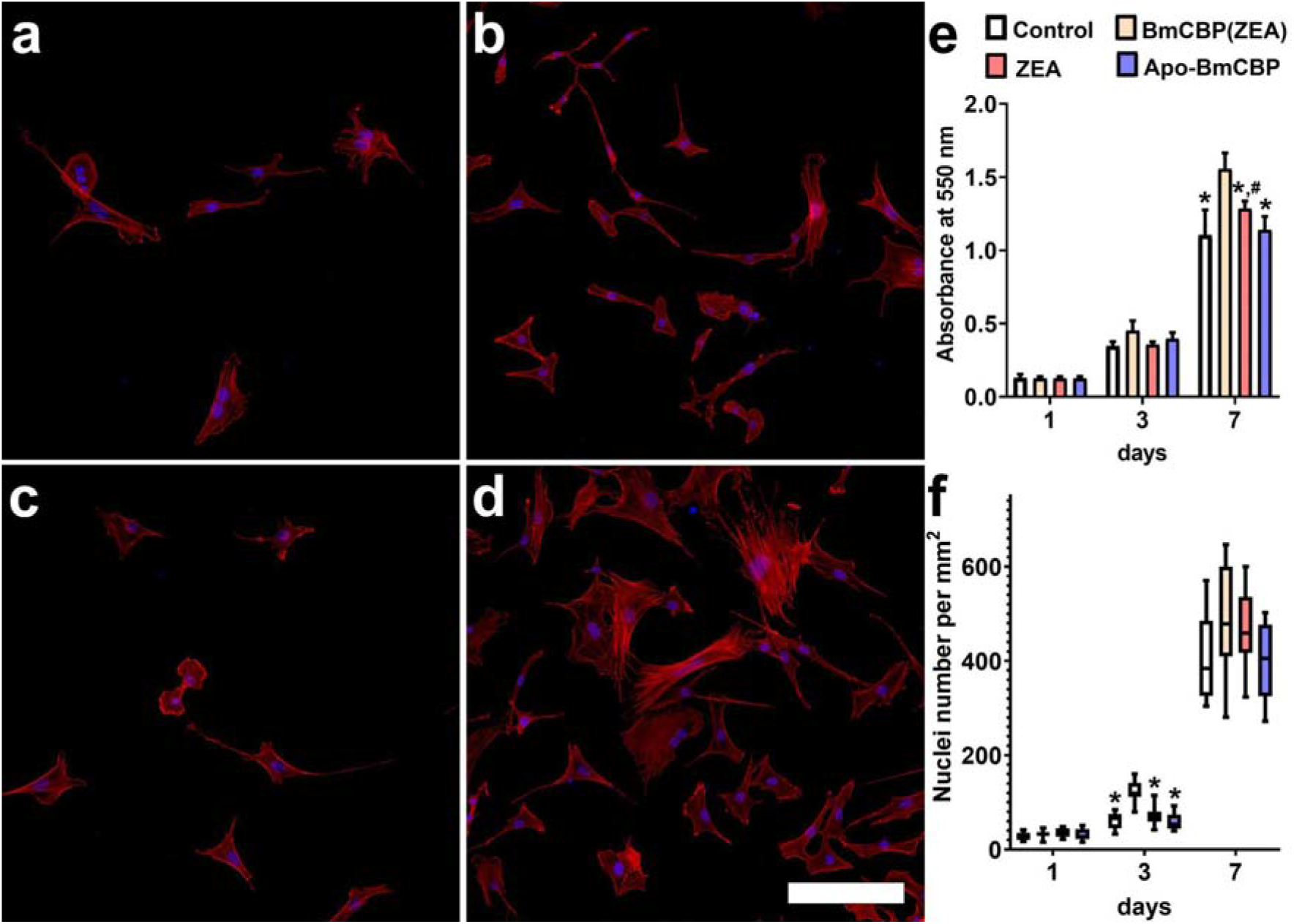
BmCBP(ZEA) stimulates mouse embryonic fibroblasts growth. a, b, c and d – 3D reconstructions made from a series of optical sections of MEFs on 1^st^ and 3^rd^ day of cultivation without (a, b) and with the addition of 0.5 μM BmCBP(ZEA) (c, d). The nuclei were visualized with Hoechst 33342 (blue); actin filaments were stained with TRITC-phalloidin (red). A scale bar equals 100 μm. e. The results of the MTT-test (mean ±SD; n=9). f. Number of cells per mm^2^ on the 1^st^, 3^rd^ and 7^th^ day of cultivation (mean ±SD; n=13). *- difference between the cells cultured with BmCBP(ZEA) and other groups. # - difference between control and other groups. p<0.05.

## Conclusions

We present here BmCBP as a universal proteinaceous nanocontainer for carotenoids and solve its crystal structure to explain the carotenoid-binding mechanism. We demonstrate that BmCBP efficiently takes up and shuttles different carotenoid types to the lipid membranes and photoactive carotenoprotein OCP – a promising optogenetic tool ^45^ and intracellular thermosensor ^39^. The carotenoids transferred by BmCBP are of paramount importance to human health ^46^, and the BmCBP-carotenoid complexes show the potential for stimulating growth of model fibroblasts. Cost-effective and easily scalable enrichment by recombinant BmCBP apoprotein of the carotenoid fraction from various crude herbal extracts is attractive for biotechnological processes. In addition, BmCBP retains its activity in modular systems (e.g., as fusion constructs) tailored for targeted delivery of lipophilic antioxidants, and thus expands the toolkit of useful water-soluble carotenoproteins. The crystal structures of BmCBP pave the way for its further bioengineering. Our complex study will serve as a blueprint for studying other START domain homologs and carotenoid-binding proteins to provide for a bigger picture.

## Methods

### Materials

All-trans-astaxanthin and β-carotene (CAS Numbers: 472-61-7 and 7235-40-7) were purchased from Sigma-Aldrich (USA). Lutein was HPLC-purified from a commercial lutein preparation purchased from RealCaps, Russia, as before ^47^. Zeaxanthin and canthaxanthin were extracted by acetone from either carotenoid-bound proteins or *E. coli* membranes as described earlier ^5,48^. Absorbance spectra of carotenoids in organic solvents were registered on a Nanophotometer NP80 (Implen, Germany) using the following molar extinction coefficients: 121,300 M^-1^ cm^-1^ for βCar at 466 nm in DMSO ^49^, 90,000 M^-1^ cm^-1^ for CAN at 472 nm in ethanol ^50^, 125,000 M^-1^ cm^-1^ for AXT at 482 nm in DMSO ^51^, 145,000 M^-1^ cm^-1^ for ZEA at 450 nm in methanol ^52^. TRITC-conjugated phalloidin and Hoechst 33342 were purchased from Thermo Fisher (USA).

Liposomes were prepared ^53^ from L-α-lecithin (20 mg) isolated from soybeans containing 17% phosphatidylcholine on a 1 ml of 50 mM KH_2_PO_4_ buffer containing 2 mM MgSO_4_, pH 7.5. The resulting mixture was homogenized in a glass homogenizer and placed in an Eppendorf tube for further sonication during 40 min on an UZDN-2T ultrasonic disintegrator (Ukrrospribor, Ukraine) at a 22 kHz frequency until the suspension became absolutely clear. Liposomes were stored at 4 °C and used within three days. Chemicals were of the highest quality and purity available.

### Cloning, protein expression and purification

cDNA corresponding to BmCBP residues 68-297 (Uniprot Q8MYA9) was codon-optimized for expression in *E. coli*, synthesized by Integrated DNA Technologies (Coralville, Iowa, USA) and cloned into the pET28b-His-3C vector (kanamycin resistance) using the *NdeI* and *XhoI* restriction sites ^25^. BmCBP mutants W232F, S206V, D162L, Q166D and I183N were obtained by the megaprimer method using Q5 (NEB) polymerase, mutagenic primers listed in Supplementary Table S3 and the WT plasmid as template. To make the design consistent, human STARD3 (residues 216-444 of Uniprot Q14849) was cloned into the same vector as BmCBP by moving from the RSFduet plasmid ^54^. The resulting constructs were verified by DNA sequencing in Evrogen (Moscow, Russia).

The wild-type BmCBP, its mutant constructs, hSTARD3 or *Synechocystis* OCP were transformed into C41(DE3) *E. coli* cells for expression of the apoforms. For reconstructing carotenoid-protein complexes, the desired plasmids were used to transform C41(DE3) cells already carrying the pACCAR25ΔcrtX plasmid (chloramphenicol resistance), which harbors the gene cluster including crtY, crtI, crtB, crtZ and crtE sequences from *Erwinia uredovora* for ZEA expression ^5,48^. Alternatively, for reconstitution of BmCBP complexes with CAN, co-expression with the ketolase crtW was used ^5^. Protein expression induced by 0.1 mM IPTG lasted overnight at 25 °C. The apo and holoforms of proteins were purified according to the unified scheme consisting of immobilized metal-affinity and size-exclusion chromatography. To improve the yield of the BmCBP holoform with ZEA, we added BSA during cell lysis and additionally used chromatography on a hydroxyapatite column. Carotenoid content of the BmCBP holoforms was verified by acetone extraction and thin-layer chromatography ^5,13^. Purified proteins were stored frozen at −80 °C.

### Circular dichroism in the visible region

His-tagged BmCBP(ZEA) (0.5 mg/ml, Vis/UV absorbance ratio of 1.4) was dialyzed overnight against 20 mM Na-phosphate buffer pH 7.1 and centrifuged for 10 min at 4 °C and 14,200g before measurements. Visible-CD spectra were recorded at 20 °C in the range of 250-650 nm at a rate of 0.4 nm/s with 1.0 nm steps in 0.1 cm quartz cuvette on a Chirascan circular dichroism spectrometer (Applied Photophysics) equipped with a temperature controller, and then buffer-subtracted. As a control, free ZEA (10 μM) in methanol was measured. ZEA concentration in methanol was determined spectrophotometrically.

### Raman spectroscopy

Raman spectroscopy was used to study the conformation of carotenoids in proteins and organic solvent. The Ntegra Spectra confocal microscope (NTMDT, Russia) equipped with a CCD spectrometer was used for examination of both liquid and solid samples. Linearly polarized light from the 532-nm laser was focused on the object using 20X (NA=0.4) or 50X (NA=0.8) Olympus (Japan) lenses. Liquid samples were placed under a microscope in quartz capillaries. The average laser power at the lens exit was about 0.5 mW. The accumulation time of single spectra varied from 10 s to 2 min depending on the type of measurements and the strength of the signal.

### Differential scanning calorimetry

The apoform or two ZEA-bound forms differing by the relative amount of the apoform (Vis/UV absorbance ratios of ~1.3 and ~1.5) of the His-tagged BmCBP (1.5 mg/ml) were dialyzed overnight against a 50 mM Na-phosphate buffer (pH 7.45), 150 mM NaCl and subjected to DSC on a VP-capillary DSC (Malvern) at a heating rate of 1 °C/min. Thermograms were processed using Origin Pro 8.0 and transition temperature (*T*_m_) was determined from the maximum of the thermal transition.

### Spectrochromatography

SEC with diode-array detection was used to analyze protein holoforms and products of carotenoid transfer. Samples (50 μl) were loaded on a Superdex 200 Increase column (GE Healthcare, Chicago, Illinois, USA) pre-equilibrated with a 20 mM Tris-HCl buffer, pH 7.6, containing 150 mM NaCl and operated using a Varian ProStar 335 system (Varian Inc., Melbourne, Australia). Flow rates and column sizes are specified in figure legends. During the runs, absorbance in the 240-900 nm range was recorded with 1-nm steps (4 nm slit width) and a 2.5 Hz frequency. Diode-array data were converted into .csv files using a custom-built Python script and processed into contour plots using Origin 9.0 (Originlab, Northampton, MA, USA).

### Carotenoid extraction by BmCBP apoprotein

Herbal extracts were prepared by mixing 1 g of each herb with 0.5 g of Na_2_SO_4_ and 0.5 g Na_2_CO_3_, followed by consecutive methanolic extraction cycles (5 ml 100% methanol) alternated by rounds of centrifugation, and the first supernatant was discarded. 2 μl out of 13 ml of each total extract were then mixed with 53 μl of 1 mg/ml His-tagged BmCBP apoprotein, incubated for 30 min at room temperature. After centrifugation, 50 μl of each sample thus obtained were subjected to IMAC and then analyzed by spectrochromatography with diode array detection. As a reference, we used a methanolic extract prepared from one tablet of the commercial food supplement Ocuvite (Lot. 2939FT139; Bausch and Lomb) containing 10 mg LUT, 2 mg ZEA and 300 μg lycopene, vitamins and microelements.

### Carotenoid transfer

Reconstitution of BmCBP holoforms with astaxanthin (AXT) and β-carotene (βCar) was accomplished by mixing 20 μl of 20 μM BmCBP apoprotein with either 2 μl buffer (negative control) or 2 μl of carotenoid solutions in DMSO (βCar 60 μM, CAN 62 μM, AXT 530 μM). CAN-bound BmCBP was obtained by adding CAN extract in DMSO, similar to that in the case of AXT and βCar, or by expression in CAN-producing *E. coli* cells 5.

Physical migration of carotenoids between proteins and membranes was studied by spectrochromatography. To this end, 10 μl of 24 μM (calculated by protein) carotenoprotein (BmCBP(ZEA) or MBP-BmCBP(ZEA)) were mixed with either 10 μl of SEC buffer (negative control) or 10 μl of 3.6 mg/ml liposomes in SEC buffer. Another control did not contain protein and only contained liposomes. Mixtures of liposomes and proteins were incubated at 25 °C for 15 min. Eighteen μl were loaded on a Superdex 200 Increase 5/150 column and analyzed by spectrochromatography. For SDS-PAGE analysis of the SEC fractions, the loading was increased 2.8 times.

CAN transfer between BmCBP and the apoform of *Synechocystis* OCP was followed by the absorbance changes in a low-binding 384-well plate with a transparent flat bottom using a Clariostar Plus plate reader (BMG, Germany) equipped with thermostat and absorbance spectrometer (200-1000 nm). Twenty μl of the His-tagged BmCBP(CAN) was mixed with 2 μl of 337 μM OCP(apo) and measured at 25 °C during 40 min with spectral measurements each 2 min. The product of carotenoid transfer was diluted to 100 μl by SEC buffer and transferred to a 1 cm quartz cuvette. For its photoconversion, a blue light-emitting diode (M455L3, Thorlabs, USA) with a maximum emission at 445 nm was used, while the steady-state absorbance spectrum was continuously registered. The temperature of the sample was stabilized by a Peltier-controlled cuvette holder Qpod 2e (Quantum Northwest, USA). Each experiment was repeated at least three times, and the most typical results are presented.

### Cell cultures and fluorescent microscopy

Primary mouse embryonic fibroblasts (MEF) were obtained as previously described ^55^. To determine the number of cells, MEF were plated onto coverslips in 35-mm Petri dishes (2×10^4^ cells in 2 ml of culture medium per coverslip). Then, the dishes were randomly distributed among the groups and 0.5 μM BmCBP(ZEA) (per carotenoid), ZEA in DMSO, Apo-BmCBP (per protein), and DMSO was used as control. After 1, 3 and 7 days of culture, the cells were fixed with 4% paraformaldehyde in phosphate-buffered saline (PBS), pH 7.4, for 30 min, washed three times with PBS, permeabilized in 0.1% Triton X-100/0.1% fetal bovine serum (FBS) solution in PBS for 30 min at 4°C, and washed twice with PBS/0.1% FBS. To identify actin microfilaments and nuclei, the cells were incubated with TRITC-conjugated phalloidin and Hoechst 33342 (both from Thermo Fisher Scientific, USA), respectively, and washed three times with PBS. The images were captured using an Eclipse Ti-E microscope with an A1 confocal module (Nikon Corporation, Japan) and a CFI Plan Apo VC 20×/0.75 objective.

### MTT-assay for cell metabolic activity

Cells were placed in 96-well plates (1500 cells per well). After 1, 3 and 7 days, 20 μl of 3-(4,5-dimethylthiazole-2-yl)-2,5-diphenyltetrazoliumbromide (MTT) solution (5 mg/ml in PBS) were added and incubated at 37 °C for 4 h. The formed formazan crystals were dissolved in DMSO and assessed colorimetrically at 550 nm.

### SEC-MALS

Size-exclusion chromatography coupled to multi-angle light scattering (SEC-MALS) was carried out using a Superdex 200 Increase 10/300 column (GE Healthcare) and a combination of a UV-Vis Prostar 335 (Varian, Australia) and a miniDAWN (Wyatt Technology, USA) detectors connected sequentially. The His-tagged BmCBP(ZEA) (4 mg/ml in 40 μl) or hSTARD3 (3.4 mg/ml in 25 μl) were loaded on the column equilibrated with filtered (0.1 μm) and degassed 20 mM Tris-HCl buffer, pH 7.6, containing 150 mM NaCl. Flow rate was 0.8 ml/min. Data were analyzed in ASTRA 8.0 (Wyatt Technology, USA) using dn/dc = 0.185 and protein extinction coefficients ε(0.1%) at 280 nm equal to 1.54 (BmCBP(ZEA)) and 0.97 (hSTARD3 apo).

### Crystallization of BmCBP

An initial crystallization screening of His-tagged BmCBP (WT apo), BmCBP(ZEA) and the W232F mutant was performed with a robotic crystallization system (Oryx4, Douglas Instruments, UK) and commercially available crystallization screens (Hampton Research, USA) using sitting drop vapor diffusion at 15 °C. The protein concentrations were 6.5 mg/ml in 20 mM Tris-HCl buffer pH 7.6 containing either 50 mM NaCl (WT) or 150 mM NaCl (W232F). The drop volume was 0.4 μl with a 50:50 and a 75:25 protein-to-precipitant ratio. Optimization of the initial conditions was made by hanging drop vapor diffusion in a 24-well plate with a 2 μl drop volume (50:50 ratio). Best crystals were obtained at 15 °C using crystallization conditions listed in Supplementary Table 2. The conditions optimized for the W232F mutant were used to crystallize the S206V and D162L mutants.

### Data collection, structure determination and refinement

BmCBP crystals were briefly soaked in a mother liquor containing either 1.4 M sodium citrate tribasic dihydrate (WT ZEA, S206V and W232F) or 20% glycerol (WT apo, D162L) immediately prior to diffraction data collection and flash-frozen in liquid nitrogen. The data were collected at 100 K at ID30-A3, ID23-2 beamlines (ESRF, France) and Rigaku OD XtaLAB Synergy-S diffractometer (IOC RAS, Moscow, Russia). Indexing, integration and scaling were done using XDS ^56^ and Dials ^57^.

The WT apo structure was solved by molecular replacement using MOLREP ^58^ and the hSTARD3 structure (PDB ID 5I9J) with removed loop regions as an initial model. For WT ZEA and mutant forms of the protein, structures were solved using WT apo as a starting model. The refinement was carried out using REFMAC5 ^59^ and BUSTER ^60^. The anisotropic or isotropic individual atom B-factors were used during the refinement for WT apo or all other structures, respectively. In all cases, hydrogens in riding positions as well as TLS were used during the refinement. The visual inspection of electron density maps and manual model rebuilding were carried out in COOT ^61^.

For representing surface electrostatic potentials, we used Adaptive Poisson-Boltzmann Solver (APBS) tools plug-in for PyMol and the default parameters (37 °C, 150 mM concentrations of negatively and positively charged ions with radii of 1.8 Å and 2.0 Å, respectively).

### Small-angle X-ray scattering

His-tagged BmCBP(ZEA) (60 μl, 13 mg/ml) was loaded onto a Superdex 200 Increase 3.2/300 column (Cytiva) and eluted at a 0.075 ml/min flow rate while the SAXS data (I(s) versus s, where s = 4πsinθ/λ, 2θ is the scattering angle and λ = 0.96787 Å) were measured at the BM29 beam line (ESRF, Grenoble, France) using a Pilatus 2M detector (data collection rate 0.5 frame/s; experiment session data DOI 10.15151/ESRF-ES-642726753). The buffer contained 20 mM Tris-HCl, pH 7.6, and 150 mM NaCl. SAXS frames recorded along the SEC profile were processed in CHROMIXS ^62^ to get an average SAXS curve corresponding to the BmCBP peak. The obtained SAXS profile was further used for modeling using the CORAL component of the ATSAS 2.8 package ^33^ whereby the crystallographic monomer was supplemented with 18 N-terminal residues while minimizing the discrepancy between the calculated scattering profile and the experimental data. This fitting procedure showed high convergence *X*^2^ values for ten CORAL-derived models were in the range 1.02-1.04 on the whole range of scattering vectors).

## Abbreviations used

AL: actinic light
βCar: β-carotene
ASESC: analytical size-exclusion spectrochromatography
AstaP: astaxanthin-binding protein
AXT: astaxanthin
CAN: canthaxanthin
CD: circular dichroism
CTDH: C-terminal domain homolog
DA: dark-adapted
ECN: echinenone
HOOP: hydrogen-out-of-plane
IMAC: immobilized metalaffinity chromatography
IPTG: isopropyl-β-thiogalactoside
LA: light-adapted
LED: lightemitting diode
MEF: mouse embryonic fibroblasts
OCP: orange carotenoid protein
SAXS: small-angle X-ray scattering
SDS-PAGE: sodium dodecyl sulfate-polyacrylamide gel electrophoresis
SEC: size-exclusion chromatography
SEC-MALS: size-exclusion chromatography coupled to multi-angle light scattering
START: steroidogenic acute regulatory lipid transfer protein
TRITC: Tetramethylrhodamine isothiocyanate
ZEA: zeaxanthin

## Data availability

The refined models and structure factors have been deposited in the Protein Data Bank under the accession codes 7ZTQ, 7ZVR, 7ZVQ, 7ZTR, 7ZTU. All materials are available from the corresponding author upon reasonable request.

## Acknowledgements

The authors thank Ilia Chetviorkin for the Python script for processing the diode-array data, Alexandr Ashikhmin for lutein preparation and Anton Popov for help with SAXS data collection at the BM29 beam line (ESRF, Grenoble, France; experiment session data DOI 10.15151/ESRF-ES-642726753). The study was supported by the Ministry of Science and Higher education of the Russian Federation in the framework of the Agreement no. 075-15-2021-1354 (07.10.2021). Carotenoprotein expression and purification was partly supported by the Russian Foundation for Basic Research and the German Research Foundation joint grant (no. 20-54-12018 and no. FR1276/6-1). CD measurements were done at the Shared-Access Equipment Centre “Industrial Biotechnology’’ of the Federal Research Center “Fundamentals of Biotechnology” of the Russian Academy of Sciences. SYK was supported by the Program of the Ministry of Science and Higher Education of Russia (0088-2021-0009).

## Author contributions

NNS – designed experiments and initiated the study; YBS, NAE, NNS – expressed and purified proteins; LAV - crystallized proteins; YBS, NAE, AMM, SYK, NNS – performed experiments; EYP - prepared and characterized liposomes; SYK - performed DSC experiments; MEM, NNS, KMB - collected X-ray data; NNS, KMB - solved crystal structures; NNS - performed SAXS data analysis; YBS, EGM, TF, NNS, KMB – analyzed data and discussed the results; VOP - supervised the study and acquired funding; NNS wrote the paper.

## Supplementary information

## Supplementary Tables

**Table S1.** Diffraction data collection and refinement statistics for BmCBP variants.

| Protein | WT apo | WT ZEA | S206V | W232F | D162L |
| --- | --- | --- | --- | --- | --- |
| **Data collection** |  |  |  |  |  |
| Diffraction source | ID23-2, ESRF | IOC RAS | IOC RAS | IOC RAS | ID30-A3, ESRF |
| Wavelength, (Å) | 0.87 | 1.54 | 1.54 | 1.54 | 0.98 |
| Crystal-to-detector distance (mm) | 185 | 36 | 45 | 36 | 130 |
| Rotation range per image (°) | 0.1 | 0.3 | 0.3 | 0.25 | 0.15 |
| Total rotation range (°) | 180 | 360 | 240 | 360 | 120 |
| Space group | C 2 2 2<sub>1</sub> | C 2 2 2<sub>1</sub> | C 2 2 2<sub>1</sub> | C 2 2 2<sub>1</sub> | C 2 2 2<sub>1</sub> |
| a,b,c, (Å) | 65.91, 67.33, 121.64 | 62.69, 66.53, 120.52 | 61.99, 65.33, 118.98, | 62.03, 66.77, 120.50, | 63.17, 67.11, 121.12, |
| Resolution range (Å) | 47.10-1.45 (1.48-1.45)* | 60.26-2.00 (2.05-2.0) | 21.03-2.50 (2.60-2.50) | 60.25 - 1.75 (1.78 - 1.75) | 46.00 - 1.90 (1.94 - 1.90) |
| Unique reflections | 47986 (2219) | 17115 (1207) | 8648 (946) | 25252 (1143) | 19947 (1268) |
| Completeness (%) | 99.7 (94.8) | 98.0 (95.5) | 99.8 (99.9) | 98.5 (84.8) | 97.3 (97.2) |
| Average redundancy | 6.3 (4.6) | 10.6 (10.3) | 8.2 (8.8) | 11.5 (4.8) | 4.5 (4.5) |
| I/sigma | 20.7 (1.3) | 15.0 (2.0) | 8.6 (1.9) | 20.8 (1.8) | 8.6 (1.4) |
| CC<sub>1/2</sub> (%) | 100.0 (49.7) | 83.4 (65.7) | 99.4 (84.2) | 97.9 (60.9) | 96.7 (74.3) |
| R<sub>pim</sub> (%) | 1.7 (56.9) | 4.3 (37.1) | 6.0 (32.2) | 2.7 (41.8) | 9.2 (79.9) |
| **Refinement** |  |  |  |  |  |
| Reflections in refinement | 47946 | 16731 | 8609 | 25215 | 19937 |
| R<sub>work</sub> (%) | 14.3 | 23.5 | 19.8 | 18.1 | 20.6 |
| R<sub>free</sub> (%) | 19.3 | 28.4 | 25.4 | 23.2 | 25.1 |
| RMS bonds (Å) | 0.02 | 0.01 | 0.01 | 0.02 | 0.02 |
| RMS angles (Å) | 2.20 | 1.63 | 1.63 | 2.09 | 2.16 |
| **Ramachandran plot** |  |  |  |  |  |
| Ramachandran favored (%) | 98.7 | 98.3 | 96.5 | 97.4 | 97.8 |
| Ramachandran allowed (%) | 1.3 | 1.3 | 3.5 | 1.8 | 2.2 |
| **Number of atoms** |  |  |  |  |  |
| Protein | 1945 | 1893 | 1788 | 1839 | 1837 |
| Ligands | 6 | 20 | - | - | - |
| Solvent | 215 | 136 | 47 | 179 | 132 |
| **Average B-factors (Å<sup>2</sup>)** |  |  |  |  |  |
| Protein | 29.08 | 41.60 | 40.90 | 25.47 | 37.22 |
| Ligands | 42.33 | 32.90 | - | - | - |
| Solvent | 41.55 | 42.10 | 35.40 | 31.36 | 39.51 |
| MolProbity | 1.45 | 1.69 | 1.80 | 1.39 | 1.44 |
| **PDB ID** | **7ZTQ** | **7ZVR** | **7ZVQ** | **7ZTR** | **7ZTU** |

\*Statistics for the highest-resolution shell is shown in parentheses.

**Table S2.** The optimized crystallization conditions for BmCBP (WT apo), BmCBP with ZEA (WT ZEA) and mutant apoproteins: W232F, S206V and D162L.

|  | WT apo | WT ZEA | W232F | S206V | D162L |
| --- | --- | --- | --- | --- | --- |
| Crystallization condition | 5% v/v<br>Tacsimate, pH<br>7.0, 0.1 M<br>HEPES, pH<br>7.0, 10% w/v<br>Polyethylene<br>glycol<br>monomethyl<br>ether 5000 | 0.7 M<br>Sodium<br>citrate<br>tribasic<br>dihydrate,<br>0.1 M BIS-TRIS<br>propane,<br>pH 7.0 | 0.5 M<br>Sodium<br>citrate<br>tribasic<br>dihydrate,<br>0.1 M BIS-TRIS<br>propane, pH<br>7.0 | 0.5 M<br>Sodium<br>citrate<br>tribasic<br>dihydrate,<br>0.1 M BIS-TRIS<br>propane, pH<br>7.0 | 0.7 M<br>Sodium<br>citrate<br>tribasic<br>dihydrate,<br>0.1 M BIS-TRIS<br>propane,<br>pH 7.0 |
| Cryoprotectant | 20% glycerol | 1.4 M<br>sodium<br>citrate<br>tribasic<br>dihydrate | 1.4 M<br>sodium<br>citrate<br>tribasic<br>dihydrate | 1.4 M<br>sodium<br>citrate<br>tribasic<br>dihydrate | 20%<br>glycerol |

**Table S3.** SAXS-derived structural parameters of His-tagged BmCBP(ZEA).

| Parameter | Value |
| --- | --- |
| Mw calculated from sequence | 28.5 kDa |
| Number of residues | 253 |
| Protein concentration | 13.1 mg/ml 60 μl loaded on a Superdex 200 Increase 3.2/300 column |
| *R*<sub>g</sub> (Guinier) | 2.12 ± 0.01 nm* |
| s*R*<sub>g</sub> limits | 0.59-1.30 |
| *R*<sub>g</sub> (reverse) | 2.19 ± 0.03 nm |
| *D*<sub>max</sub> | 8.5 nm |
| Porod volume | 51,057 nm<sup>3</sup> |
| *M*<sub>w</sub> Porod | 31.9 kDa |
| *M*<sub>w</sub> MoW | 31.2 kDa |
| *M*<sub>w</sub> Vc | 27.6 kDa |
| Kratky plot | bell-shaped (folded) |

\*All SAXS-derived parameters were calculated using the ATSAS 2.8 software package<sup>38</sup>.

**Table S4.**
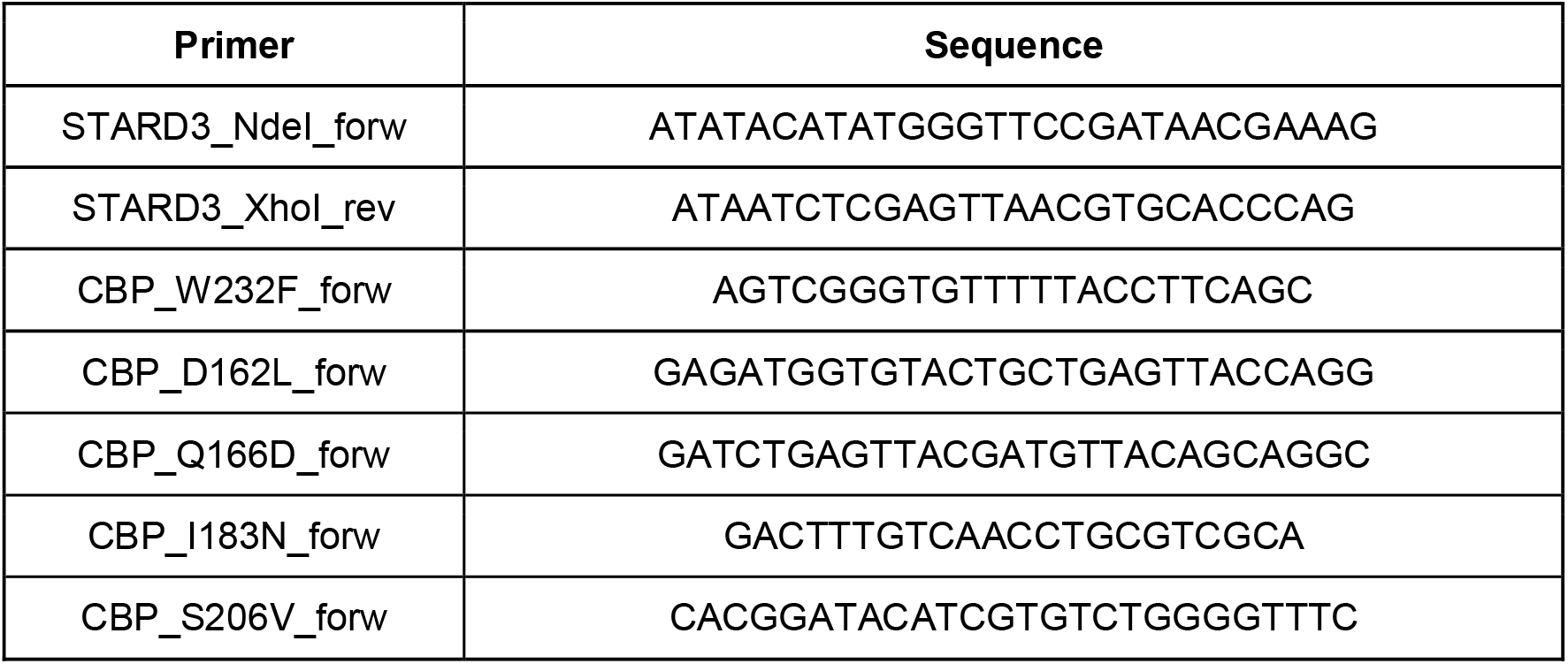
Primers used in this study.

## Supplementary figures

**Fig. S1.**
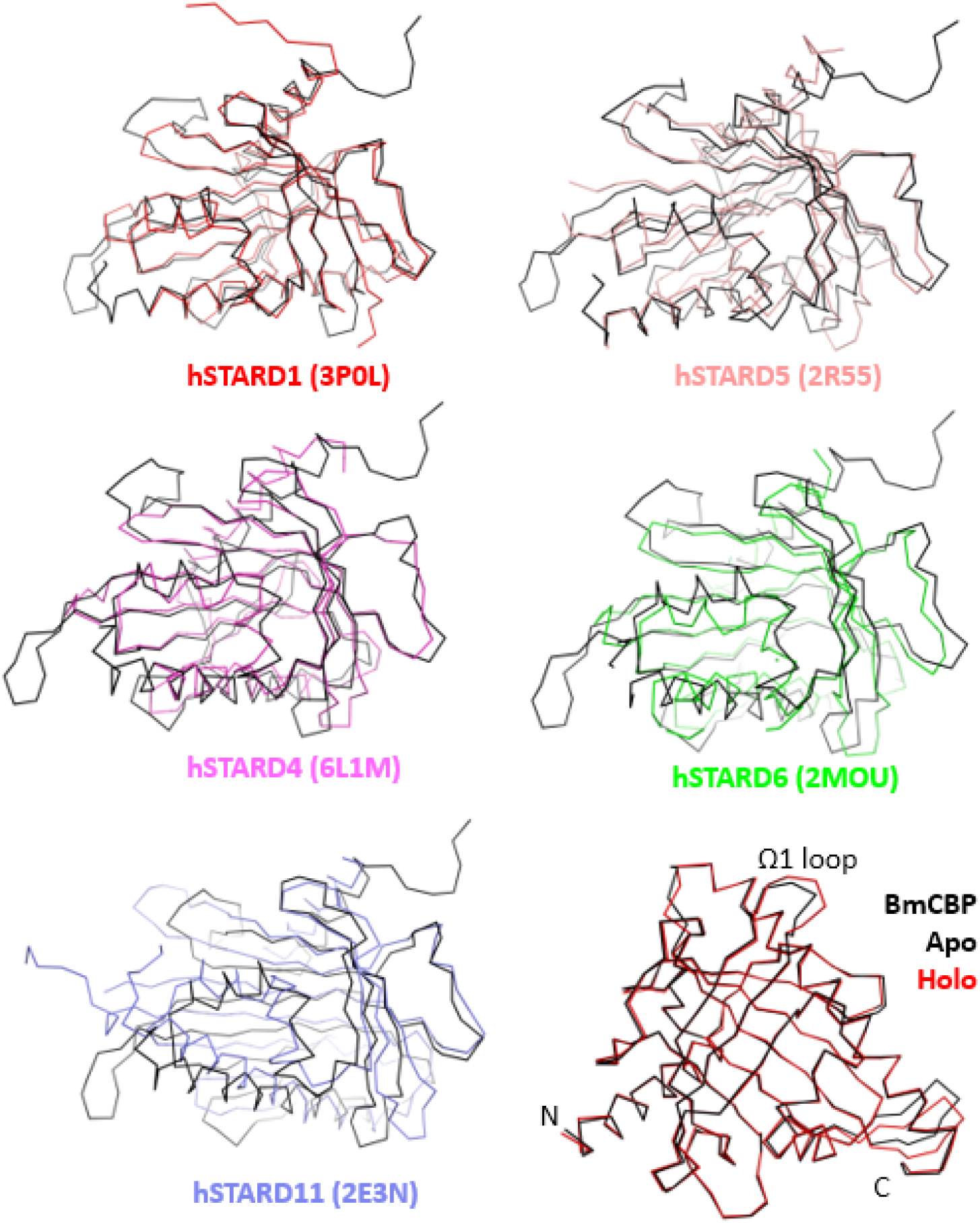
BmCBP has a START-like fold. Superposition of various human START protein domains with that of BmCBP shown as ribbon diagrams. Human STARD1, STARD4, STARD5, STARD6 and STARD11 are shown by different colors as indicated, along with their corresponding PDB IDs. BmCBP is shown in black ribbon for clarity. Bottom right corner, superimposition of the apo and ZEA-bound BmCBP structures solved in this work.

**Fig. S2.**
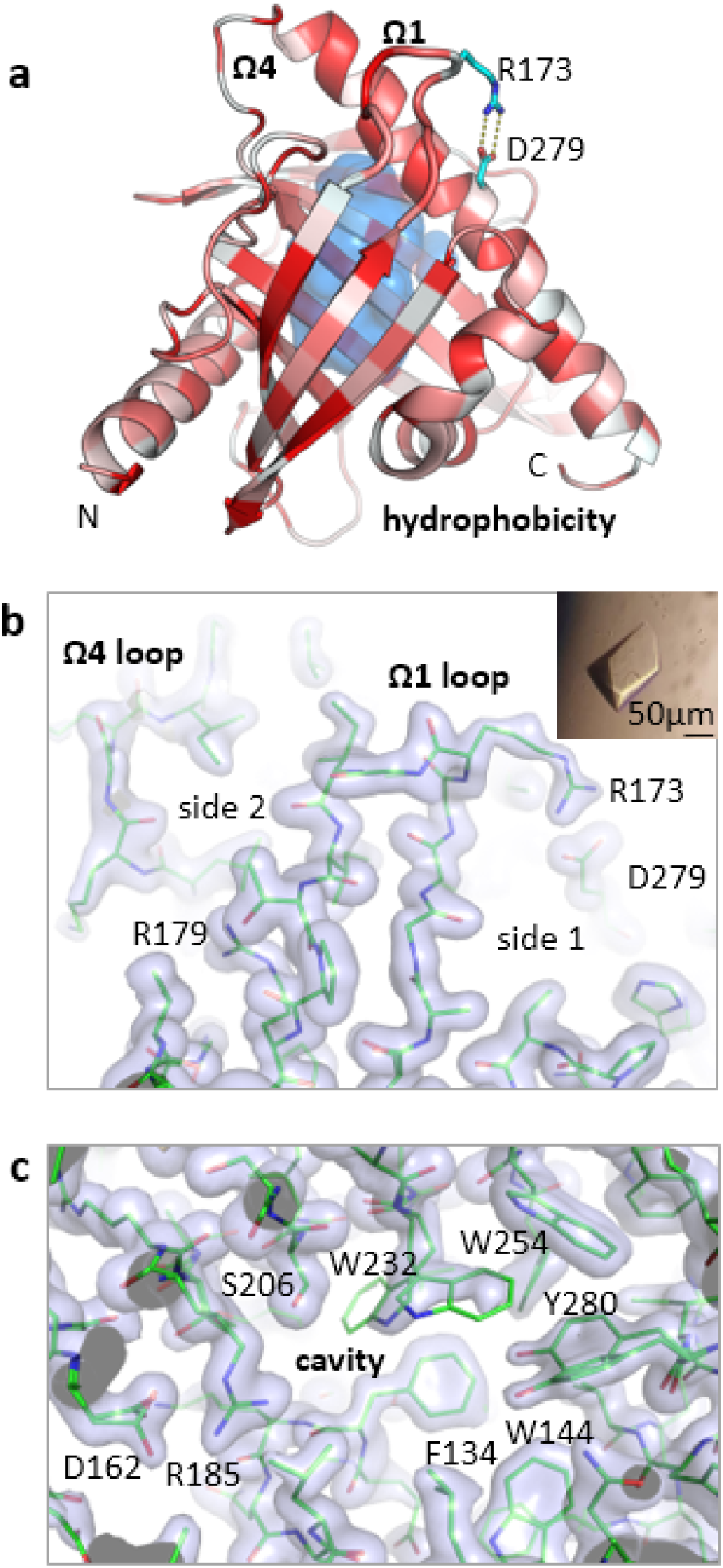
Crystal structure of BmCBP apoprotein. a. BmCBP structure is colored according to the Eisenberg hydrophobicity scale (red - hydrophobic, white - polar) ^63^. The ligand-binding cavity is shown by an internal semi-transparent blue surface. The unique salt bridge between Arg173 and Asp279 attaching the Ω1-loop to the □4 helix is shown. b. A fragment of the 2Fo-Fc electron density map (1σ) showing contacts of the Ω1-loop from the side of the salt bridge (side 1) and the hydrophobic side (side 2). The gatekeeper Arg179 at the base of the Ω1 loop is labeled. The insert shows the appearance of the crystal. c. A fragment of the 2Fo-Fc map (1σ) showing the BmCBP interior with several residues having alternative conformations.

**Fig. S3.**
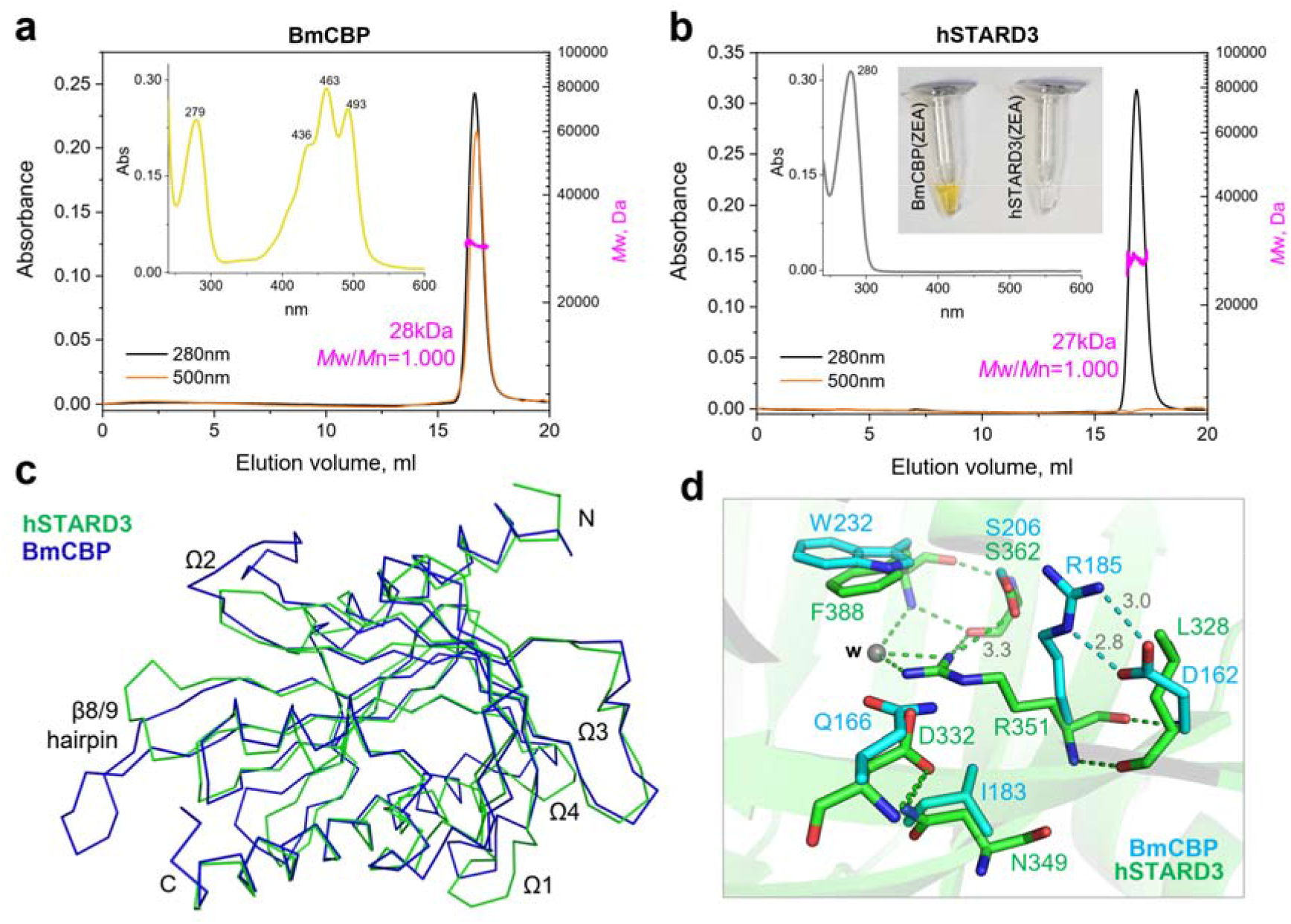
Unlike BmCBP, hSTARD3 does not bind ZEA. a, b. Analysis of BmCBP (a) or hSTARD3 (b) ability to mature into holoforms upon expression in ZEA-synthesizing *E.coli* cells. The His-tagged proteins were purified by IMAC and then analyzed by spectrochromatography (Superdex 200 Increase 10/300, 0.8 ml/min) coupled to MALS. The inserts show absorbance spectra corresponding to the peaks on the elution profiles and the color of the samples obtained. Note the successful formation of the holoform in the case of BmCBP only. *M*_w_ distributions across the chromatography peaks are shown along with the average *M*_w_ values and polydispersity indices (*M*_w_/*M*_n_). c. Overlaid backbones of BmCBP (apo) and hSTARD3 (apo; PDB 5I9J) shown as ribbon diagrams. d. Superposition of the tentative carotenoid-binding sites of BmCBP and hSTARD3 showing key differences. Relevant distances are indicated in Å. Having the markedly different carotenoid-binding capacity, BmCBP and hSTARD3 differ by the length and/or conformation of their Ω1, Ω2 and Ω4-loops, and the β8/9 hairpin. However, the inability of hSTARD3 to yield holoforms was likely due to differences in the carotenoid-binding site. Both proteins have in place the serine (206/362) and arginine (185/351) residues that face the carotenoid ring in the BmCBP(ZEA) structure, whereas their other proximal residues are different (and also vary among the homologs, see main text): Trp232 in BmCBP is replaced by Phe388 in hSTARD3, Asp162 is replaced by Leu328, Gln166 is replaced by Asp332, and Ile183 is replaced by Asn349. In addition, we noticed that the conserved arginine (185/351) adopts different conformations in BmCBP and in hSTARD3. In BmCBP, Arg185 forms the salt bridge with Asp162, whereas in hSTARD3 the Asp162/Leu328 substitution releases the Arg351’s fall into the ligand-binding cavity, which is favored by H-bonding interactions with Ser362.

**Fig. S4.**
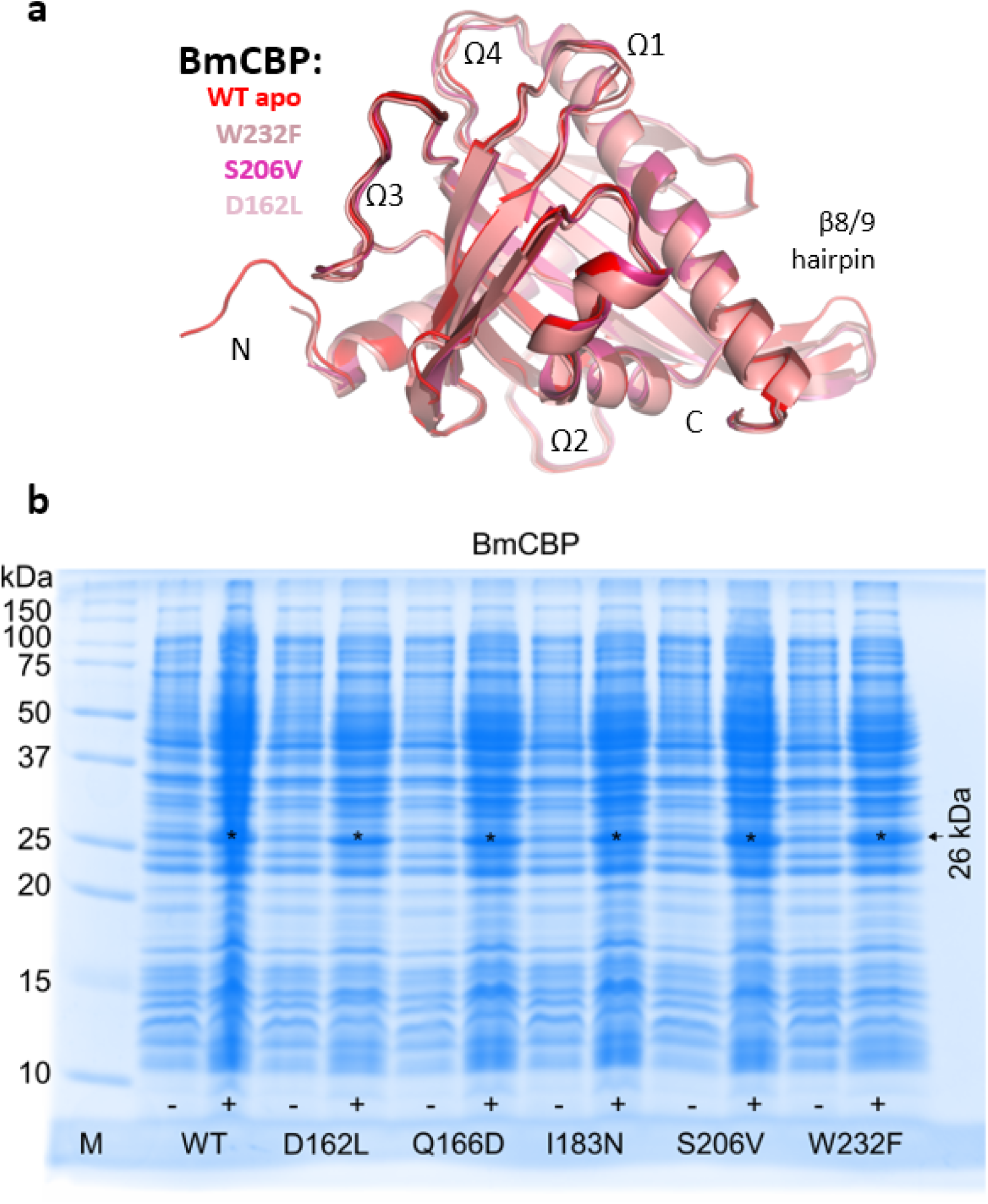
BmCBP point mutants. a, Superimposition of crystal structures of BmCBP apoproteins: WT, W232F, S206V and D162L, with the main structural elements labeled. b, SDS-PAGE analysis of expression of BmCBP WT and its mutants in ZEA-producing *E.coli* cells.

**Fig. S5.**
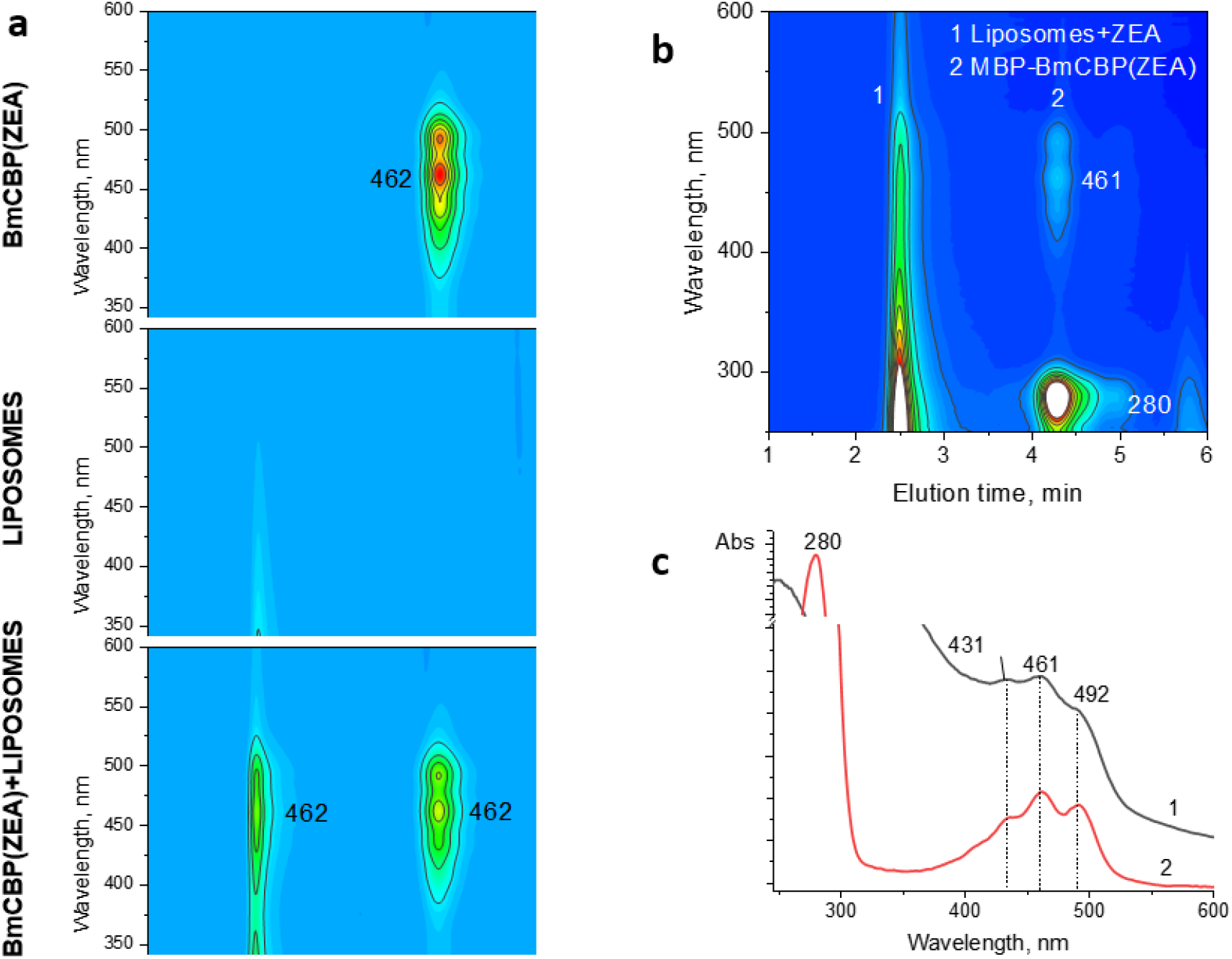
ZEA transfer from BmCBP to liposomes analyzed by spectrochromatography. a, Spectrochromatograms of BmCBP(ZEA), liposomes, or their mixture. Main absorbance maxima are indicated in nm. The samples were analyzed after completion of the ZEA transfer process. b, BmCBP retains the ability to load and transfer ZEA to liposomes even when fused to a bigger maltose-binding protein (MBP). A spectrochromatogram showing the result of the transfer (b, raw data) and ZEA redistribution between the liposome and protein fractions (c). 1 and 2 fractions are named in the legend.

## References

1. Cianci, M. et al. The molecular basis of the coloration mechanism in lobster shell: beta-crustacyanin at 3.2-A resolution. Proc Natl Acad Sci U A 99, 9795–800 (2002).

2. Chayen, N. E. et al. Unravelling the structural chemistry of the colouration mechanism in lobster shell. Acta Crystallogr Biol Crystallogr 59, 2072–82 (2003).

3. Hara, K. Y., Yagi, S., Hirono-Hara, Y. & Kikukawa, H. A Method of Solubilizing and Concentrating Astaxanthin and Other Carotenoids. Mar. Drugs 19, 462 (2021).

4. Kawasaki, S., Mizuguchi, K., Sato, M., Kono, T. & Shimizu, H. A novel astaxanthin-binding photooxidative stress-inducible aqueous carotenoprotein from a eukaryotic microalga isolated from asphalt in midsummer. Plant Cell Physiol 54, 1027–40 (2013).

5. Slonimskiy, Y. B., Egorkin, N. A., Friedrich, T., Maksimov, E. G. & Sluchanko, N. N. Microalgal protein AstaP is a potent carotenoid solubilizer and delivery module with a broad carotenoid binding repertoire. FEBS J. 289, 999–1022 (2022).

6. Muzzopappa, F. & Kirilovsky, D. Changing Color for Photoprotection: The Orange Carotenoid Protein. Trends Plant Sci (2019) doi:10.1016/j.tplants.2019.09.013.

7. Wilson, A. et al. A photoactive carotenoid protein acting as light intensity sensor. Proc Natl Acad Sci U A 105, 12075–80 (2008).

8. Sedoud, A. et al. The Cyanobacterial Photoactive Orange Carotenoid Protein Is an Excellent Singlet Oxygen Quencher. Plant Cell 26, 1781–1791 (2014).

9. Kerfeld, C. A. et al. The crystal structure of a cyanobacterial water-soluble carotenoid binding protein. Structure 11, 55–65 (2003).

10. Maksimov, E. G. et al. The Unique Protein-to-Protein Carotenoid Transfer Mechanism. Biophys J 113, 402–414 (2017).

11. Moldenhauer, M. et al. Assembly of photoactive orange carotenoid protein from its domains unravels a carotenoid shuttle mechanism. Photosynth Res 133, 327–341 (2017).

12. Melnicki, M. R. et al. Structure, Diversity, and Evolution of a New Family of Soluble Carotenoid-Binding Proteins in Cyanobacteria. Mol Plant 9, 1379–1394 (2016).

13. Slonimskiy, Y. B. et al. Light-controlled carotenoid transfer between water-soluble proteins related to cyanobacterial photoprotection. FEBS J 286, 1908–1924 (2019).

14. Muzzopappa, F. et al. Paralogs of the C-Terminal Domain of the Cyanobacterial Orange Carotenoid Protein Are Carotenoid Donors to Helical Carotenoid Proteins. Plant Physiol 175, 1283–1303 (2017).

15. Maksimov, E. G. et al. Soluble Cyanobacterial Carotenoprotein as a Robust Antioxidant Nanocarrier and Delivery Module. Antioxid. Basel 9, (2020).

16. Harris, D. et al. Structural rearrangements in the C-terminal domain homolog of Orange Carotenoid Protein are crucial for carotenoid transfer. Commun Biol 1, 125 (2018).

17. Alpy, F. & Tomasetto, C. Give lipids a START: the StAR-related lipid transfer (START) domain in mammals. J Cell Sci 118, 2791–801 (2005).

18. Li, B., Vachali, P., Frederick, J. M. & Bernstein, P. S. Identification of StARD3 as a lutein-binding protein in the macula of the primate retina. Biochemistry 50, 2541–9 (2011).

19. Arunkumar, R., Gorusupudi, A. & Bernstein, P. S. The macular carotenoids: A biochemical overview. Biochim. Biophys. Acta BBA - Mol. Cell Biol. Lipids 1865, 158617 (2020).

20. Moeller, S. M., Jacques, P. F. & Blumberg, J. B. The Potential Role of Dietary Xanthophylls in Cataract and Age-Related Macular Degeneration. J. Am. Coll. Nutr. 19, 522S–527S (2000).

21. Horvath, M. P. et al. Structure of the lutein-binding domain of human StARD3 at 1.74 A resolution and model of a complex with lutein. Acta Crystallogr F Struct Biol Commun 72, 609–18 (2016).

22. Tsujishita, Y. & Hurley, J. H. Structure and lipid transport mechanism of a StAR-related domain. Nat Struct Biol 7, 408–14 (2000).

23. Tabunoki, H. et al. Isolation, characterization, and cDNA sequence of a carotenoid binding protein from the silk gland of Bombyx mori larvae. J Biol Chem 277, 32133–40 (2002).

24. Sakudoh, T. et al. Carotenoid silk coloration is controlled by a carotenoid-binding protein, a product of the Yellow blood gene. Proc. Natl. Acad. Sci. 104, 8941–8946 (2007).

25. Slonimskiy, Y. B. et al. Reconstitution of the functional carotenoid-binding protein from silkworm in E. coli. Int. J. Biol. Macromol. (2022) doi:10.1016/j.ijbiomac.2022.06.135.

26. Murcia, M., Faraldo-Gomez, J. D., Maxfield, F. R. & Roux, B. Modeling the structure of the StART domains of MLN64 and StAR proteins in complex with cholesterol. J Lipid Res 47, 2614–30 (2006).

27. Letourneau, D. et al. STARD6 on steroids: solution structure, multiple timescale backbone dynamics and ligand binding mechanism. Sci Rep 6, 28486 (2016).

28. Sluchanko, N. N., Tugaeva, K. V. & Maksimov, E. G. Solution structure of human steroidogenic acute regulatory protein STARD1 studied by small-angle X-ray scattering. Biochem Biophys Res Commun 489, 445–450 (2017).

29. Chovancova, E. et al. CAVER 3.0: A Tool for the Analysis of Transport Pathways in Dynamic Protein Structures. PLOS Comput. Biol. 8, e1002708 (2012).

30. Liebschner, D. et al. Polder maps: improving OMIT maps by excluding bulk solvent. Acta Crystallogr. Sect. Struct. Biol. 73, 148–157 (2017).

31. Carotenoids, Volume 1B: Spectroscopy.

32. de Faria, A. F., de Rosso, V. V. & Mercadante, A. Z. Carotenoid Composition of Jackfruit (Artocarpus heterophyllus), Determined by HPLC-PDA-MS/MS. Plant Foods Hum. Nutr. 64, 108–115 (2009).

33. Petoukhov, M. V. et al. New developments in the ATSAS program package for small-angle scattering data analysis. J Appl Cryst 45, 342–350 (2012).

34. Xue, Y. et al. Identification of a key gene StAR-like-3 responsible for carotenoids accumulation in the noble scallop Chlamys nobilis. Food Chem. Mol. Sci. 4, 100072 (2022).

35. Pishchalnikov, R. Y. et al. Structural peculiarities of keto-carotenoids in water-soluble proteins revealed by simulation of linear absorption. Phys Chem Chem Phys 21, 25707–25719 (2019).

36. Maksimov, E. G. et al. The Signaling State of Orange Carotenoid Protein. Biophys J 109, 595–607 (2015).

37. Leverenz, R. L. et al. Structural and functional modularity of the orange carotenoid protein: distinct roles for the N-and C-terminal domains in cyanobacterial photoprotection. Plant Cell 26, 426–37 (2014).

38. Dudek, M. et al. Chiral Amplification in Nature: Studying Cell-Extracted Chiral Carotenoid Microcrystals via the Resonance Raman Optical Activity of Model Systems. Angew. Chem. Int. Ed. 58, 8383–8388 (2019).

39. Maksimov, E. G. et al. A genetically encoded fluorescent temperature sensor derived from the photoactive Orange Carotenoid Protein. Sci Rep 9, 8937 (2019).

40. Gurunathan, S. et al. Cytotoxicity and Transcriptomic Analysis of Silver Nanoparticles in Mouse Embryonic Fibroblast Cells. Int. J. Mol. Sci. 19, 3618 (2018).

41. Harrison, E. H. Carotenoids, ß-Apocarotenoids, and Retinoids: The Long and the Short of It. Nutrients 14, 1411 (2022).

42. Szymanski, Ł. et al. Retinoic Acid and Its Derivatives in Skin. Cells 9, 2660 (2020).

43. Lenz, M. et al. All-trans retinoic acid induces synaptic plasticity in human cortical neurons. eLife 10, e63026 (2021).

44. Tang, X.-H. et al. A Retinoic Acid Receptor β2 Agonist Improves Cardiac Function in a Heart Failure Model. J. Pharmacol. Exp. Ther. (2021) doi:10.1124/jpet.121.000806.

45. Piccinini, L. et al. A synthetic switch based on orange carotenoid protein to control blue–green light responses in chloroplasts. Plant Physiol. kiac122 (2022) doi:10.1093/plphys/kiac122.

46. Mrowicka, M., Mrowicki, J., Kucharska, E. & Majsterek, I. Lutein and Zeaxanthin and Their Roles in Age-Related Macular Degeneration—Neurodegenerative Disease. Nutrients 14, 827 (2022).

47. Ashikhmin, A., Makhneva, Z., Bolshakov, M. & Moskalenko, A. Incorporation of spheroidene and spheroidenone into light-harvesting complexes from purple sulfur bacteria. J. Photochem. Photobiol. B 170, 99–107 (2017).

48. Maksimov, E. G. et al. A comparative study of three signaling forms of the orange carotenoid protein. Photosynth Res 130, 389–401 (2016).

49. Craft, N. E. & Soares, J. H. Relative solubility, stability, and absorptivity of lutein and .beta.-carotene in organic solvents. J. Agric. Food Chem. 40, 431–434 (1992).

50. Krajewska, M., Szymczak-Żyła, M. & Kowalewska, G. Carotenoid determination in recent marine sediments - practical problems during sample preparation and HPLC analysis. Curr. Chem. Lett. 91–104 (2017) doi:10.5267/j.ccl.2017.4.003.

51. Régnier, P. et al. Astaxanthin from Haematococcus pluvialis Prevents Oxidative Stress on Human Endothelial Cells without Toxicity. Mar. Drugs 13, 2857–2874 (2015).

52. Zang, L.-Y., Sommerburg, O. & van Kuijk, F. J. G. M. Absorbance Changes of Carotenoids in Different Solvents. Free Radic. Biol. Med. 23, 1086–1089 (1997).

53. Racker, E. A new procedure for the reconstitution of biologically active phospholipid vesicles. Biochem Biophys Res Commun 55, 224–30 (1973).

54. Tugaeva, K. V. et al. Molecular basis for the recognition of steroidogenic acute regulatory protein by the 14-3-3 protein family. FEBS J 287, 3944–3966 (2020).

55. Conner, D. A. Mouse Embryo Fibroblast (MEF) Feeder Cell Preparation. Curr. Protoc. Mol. Biol. 51, 23.2.1–23.2.7 (2000).

56. Kabsch, W. XDS. Acta Crystallogr. D Biol. Crystallogr. 66, 125–132 (2010).

57. Winter, G. et al. DIALS: implementation and evaluation of a new integration package. Acta Crystallogr. Sect. Struct. Biol. 74, 85–97 (2018).

58. Vagin, A. & Teplyakov, A. Molecular replacement with MOLREP. Acta Crystallogr Biol Crystallogr 66, 22–5 (2010).

59. Murshudov, G. N. et al. REFMAC5 for the refinement of macromolecular crystal structures. Acta Crystallogr. D Biol. Crystallogr. 67, 355–367 (2011).

60. Blanc, E. et al. Refinement of severely incomplete structures with maximum likelihood in BUSTER-TNT. Acta Crystallogr Biol Crystallogr 60, 2210–21 (2004).

61. Emsley, P. & Cowtan, K. Coot: model-building tools for molecular graphics. Acta Crystallogr Biol Crystallogr 60, 2126–32 (2004).

62. Panjkovich, A. & Svergun, D. I. CHROMIXS: automatic and interactive analysis of chromatography-coupled small-angle X-ray scattering data. Bioinformatics 34, 1944–1946 (2018).

63. Eisenberg, D., Schwarz, E., Komaromy, M. & Wall, R. Analysis of membrane and surface protein sequences with the hydrophobic moment plot. J Mol Biol 179, 125–42 (1984).

